# Antagonism between regular and atypical Cxcr3 receptors regulates macrophage migration during infection and injury in zebrafish

**DOI:** 10.1101/719526

**Authors:** Frida Sommer, Vincenzo Torraca, Sarah Kamel, Amber Lombardi, Annemarie H. Meijer

## Abstract

The CXCR3-CXCL11 chemokine-signaling axis plays an essential role in infection and inflammation by orchestrating leukocyte trafficking in human and animal models, including zebrafish. Atypical chemokine receptors (ACKRs) play a fundamental regulatory function in signaling networks by shaping chemokine gradients through their ligand scavenging function, while being unable to signal in the classic G-protein-dependent manner. Two copies of the *CXCR3* gene in zebrafish, *cxcr3.2 and cxcr3.3,* are expressed on macrophages and share a highly conserved ligand-binding site. However, Cxcr3.3 has structural characteristics of ACKRs indicative of a ligand-scavenging role. In contrast, we previously showed that Cxcr3.2 is an active CXCR3 receptor since it is required for macrophage motility and recruitment to sites of mycobacterial infection. In this study, we generated a *cxcr3.3* CRISPR-mutant to functionally dissect the antagonistic interplay between the *cxcr3* paralogs in the immune response. We observed that *cxcr3.3* mutants are more susceptible to mycobacterial infection, while *cxcr3.2* mutants are more resistant. Furthermore, macrophages in the *cxcr3.3* mutant are more motile, show higher activation status, and are recruited more efficiently to sites of infection or injury. Our results suggest that Cxcr3.3 is an ACKR that regulates the activity of Cxcr3.2 by scavenging common ligands and that silencing the scavenging function of Cxcr3.3 results in an exacerbated Cxcr3.2 signaling. In human, splice variants of CXCR3 have antagonistic functions and CXCR3 ligands also interact with ACKRs. Therefore, in zebrafish, an analogous regulatory mechanism appears to have evolved after the *cxcr3* gene duplication event, through diversification of conventional and atypical receptor variants.

**Summary sentence:** CXCR3 paralogue with structural characteristics of atypical chemokine receptors regulates the activity of a conventional receptor involved in macrophage motility by scavenging shared ligands.

## Introduction

Chemokine signaling is essential for the proper functioning of the immune system. Leukocyte populations differentially express chemokine receptors that participate in processes such as development, differentiation, cell proliferation, leukocyte trafficking, and immune responses [1, 2, 3, 4]. Chemokine receptors are a type of G-protein-coupled receptors (GPCRs) that belong to the class A (rhodopsin-like) family. They have the prototypal GPCR structure consisting of an extracellular NH2 terminus, an intercellular COOH terminus, and 7 transmembrane domains (TM) interconnected by 3 extracellular (EC) and 3 intracellular (IC) loops [5, 6]. This receptor class has been divided into 5 subclasses based on the pattern of highly conserved cysteine residues they display (C, CC, CXC, CX3C and, XC) and on the chemokines that they bind (CCL, CXCL, XCL, CX3CL) [6, 7]. A distinctive feature of chemokine signaling is its pleiotropic nature. Most chemokine receptors can bind multiple chemokines, and chemokines can also bind to numerous receptors [5, 2]. The redundancy of the interactions and the diversity of processes involving chemokine receptors require tightly regulated mechanisms to confer specificity to the response resulting from a receptor-ligand interaction [8, 6, 9]. Therefore, chemokine signaling-axes regulation and signal integration occur at different levels (genetic, functional, spatial and temporal) and engage a wide variety of mechanisms to evoke specific responses [10, 11, 12].

One kind of mechanism for regulating chemokine receptor activities involves atypical chemokine receptors (ACKRs), a heterogeneous group of proteins [13, 14]. Despite their structural diversity and distant evolutionary relationships, all ACKRs are unified by their inability to signal in the classic G-protein-dependent fashion and by their shared capacity to shape chemokine gradients [13, 15]. These receptors display characteristic features such as amino acid substitutions within the central activation E/DRY-motif (aspartic/glutamic acid- arginine- tyrosine- motif) [13, 16], which is crucial for G-protein coupling and further downstream signaling [16]. The central Arginine (R) of the E/DRY-motif is highly conserved (96%) among functional GPCRs as it is critical for locking and unlocking the receptor and substitutions of this residue usually result in loss of function [17, 16]. In addition, ACKRs show alterations in amino acid residues within the TM domains that function as microswitches by stabilizing the active conformation of a GPCR. ACKRs have been shown to exert their function by scavenging or sequestering chemokines or by altering the activity or membrane expression of conventional chemokine receptors [10, 13]. The functional read-out of ACKRs is that they fail to induce cell migration, contrary to the well-characterized chemotactic function of conventional chemokine receptors [13, 18].

The zebrafish model has been successfully used to functionally unravel mechanistic processes underlying chemokine networks involving ACKRs [19, 20]. The optical transparency of larvae facilitates live visualization of immunological processes and provides a reasonably simplified *in-vivo* model for chemokine signaling if used before adaptive immunity arises [21, 22, 23, 24]. Besides, due to the extensive duplication of chemokine receptor genes in teleost fish, the zebrafish provides a useful experimental system to address sub-functionalization or loss of function events. The sub-functionalization of two *CXCR4* genes, *cxcr4a* and *cxcr4b*, was determined using the zebrafish model. In several studies, *cxcr4a* was associated primarily with cell proliferation [19, 11], whereas *cxcr4b* was related to the retention of hematopoietic stem cells in hematopoietic tissue, recruitment of leukocytes to sites of infection and damage, modulation of inflammation, neutrophil migration, primordial cell and tissue migration, and tissue regeneration [25]. Cxcr4b interacts with Cxcl12a and it was shown that this chemokine is also a ligand for the scavenger receptor Cxcr7 (ACKR3) [26, 27]. Interacting with both receptors, Cxcl12a has been shown to control the migration of a tissue primordium, in which expression of *cxcr4b* and *cxcr7* is spatially restricted to the leading and trailing edge, respectively [19, 11]. The scavenging role of CXCR7 (ACKR3) in the regulation of the CXCL12-CXCR4 axis was later confirmed in human cells [26]. Moreover, the zebrafish model allowed to visualize the contribution of endogenous chemokine receptors in shaping self-generated gradients of migrating cells [20], and revealed how the cell-type expressing a given chemokine receptor is the major determinant for the functional specificity of a chemokine receptor-ligand interaction, and not the receptor-ligand pair itself [28].

The human CXCR3 chemokine receptor and its ligands (CXCL9-11) have been proven instrumental for T-cell functioning as well as for macrophage recruitment to sites of infection and injury, and are therefore implicated in several infectious and pathological conditions, including tuberculosis [29, 30]. CXCR3 ligands have been proposed as clinical markers for the diagnosis of this infectious disease and the response to treatment [31, 32]. In a previous study, we assessed the role of CXCR3 in mycobacterial infection using the zebrafish-*Mycobacterium marinum* model and observed that CXCR3 ligands were induced upon infection in this model, like in human patients [29, 33]. *Mycobacterium marinum* is a close relative of *Mycobacterium tuberculosis* and a natural pathogen of various ectotherms, such as zebrafish, which has become widely used to unravel early innate immune responses against mycobacterial infections [21, 33, 34]. In zebrafish there are three copies of the CXCR3 gene: *cxcr3.1*, *cxcr3.2*, and *cxc3.3*. We determined that the latter two are expressed on macrophages at early developmental stages as well as at 5 and 6 days post-fertilization (dpf) [35] and that *cxcr3.2* is a functional homolog of human *CXCR3* [29].

Macrophages play a pivotal role in mycobacterial infections since they are motile and phagocytic cells as well as a constituent cell-type of the characteristic granulomas that represent inflammatory infection foci [30, 33]. The efferocytosis of infected macrophages in granulomas contributes to the amplification of the infection and is a crucial process to consider to design new therapeutic strategies [21, 29]. In a previous study, we showed that Cxcr3.2 is required for the proper migration of macrophages to infectious foci [29]. However, in agreement with studies in *cxcr3* mutant mice, mutation of *cxcr3.2* is beneficial to the host in the context of mycobacterial infection [30]. We showed that *cxcr3.2* mutation favors bacterial contention, since it results in a reduced macrophage motility, thereby preventing macrophage-mediated dissemination of bacteria and limiting the expansion of granulomas.

While Cxcr3.2 is required for macrophage migration in zebrafish, the function of its paralog, Cxcr3.3, which is also expressed on macrophages, remains unknown. In the present study, we investigated the regulatory interplay between Cxcr3.2 and Cxcr3.3 in the context of *M. marinum* infection and in the response to injury, using a tail-amputation model. Opposite to *cxcr3.2* mutants, functional assays showed that *cxcr3.3* mutation leads to poor control of the infection and that *cxcr3.3* mutant macrophages are more motile and, consequently, display an enhanced recruitment to sites of infection and damage. As a result of an enhanced macrophage recruitment and an increased cell motility, bacterial dissemination is facilitated in the *cxcr3.3* mutants. Structural predictions suggest that the Cxcr3.3 receptor can bind the same ligands as Cxcr3.*2* because of the high conservation of the ligand-binding sites, but also that it cannot signal using classic G-protein-dependent pathways. Taking both our structural and functional data together, we posit that the two CXCR3 zebrafish paralogs *cxcr3.2* and *cxcr3.3* function antagonistically. We propose that Cxcr3.3 is an ACKR that functionally regulates the activity of Cxcr3.2 by scavenging common ligands and that knocking out *cxcr3.3* results in an exacerbated Cxcr3.2 signaling due to an excess of available chemokines.

## Methods

### Zebrafish lines and husbandry

Zebrafish husbandry and experiments were conducted in compliance with guidelines from the Zebrafish Model Organism Database (http://zfin.org), the EU Animal Protection Directive 2010/63/EU, and the directives of the local animal welfare committee of Leiden University (License number: 10612). All WT, mutant and transgenic lines used in this study were generated in the AB/TL background. The zebrafish lines used were: WT-AB/TL, homozygous mutant (*cxcr3.2-/-*) and WT siblings (*cxcr3.2+/+*) of *cxcr3.2^hu6044^*, homozygous mutant (*cxcr3.3-/-*) and WT siblings (*cxcr3.3+/+)* of *cxcr3.3^ibl50^,* and the same lines crossed into *Tg(mpeg1: mCherry-F)^ump2^* background and *Tg (mpx: eGFP)^i114^* [36], and homozygous mutants (*dram1-/-*)and wild type siblings (*dram1+/+*) of *dram1^ibl53^* [37]. Eggs and larvae were kept at 28.5°C in egg water (60 µg/ml Instant Ocean sea salts and 0.0025% methylene blue). All larvae were anesthetized with 0.02% buffered tricaine, (3-aminobenzoic acid ethyl ester; Sigma Aldrich, St. Louis, MO, USA) before infection, tail-amputation, and imaging. Larvae were kept in egg water containing 0.003% PTU (1-phenyl-2-thiourea, Sigma Aldrich) to prevent pigmentation before confocal imaging.

### Generation and characterization of the *cxcr3.3* mutant zebrafish line

A *cxcr3.3-/-* (*cxcr3.3^ibl50^*) zebrafish line was generated using CRISPR-Cas9 technology. Short guide RNAs (sgRNAs) targeting the proximal region of the *cxcr3.3* gene (ENSDARG00000070669) were designed using the chop-chop web-server [38, 39]. The 122 bp DNA template was generated by annealing and amplifying semi-complementary oligonucleotides using the following PCR program: initial denaturation 3 min at 95°C, 5 denaturation cycles at 95°C for 30 s, annealing for 60 s at 55°C, elongation phase for 30 s at 72°C and final extension step at 72°C for 15 min. The reaction volume was 50 µL, 200uM dNTPs and 1 unit of Dream Taq polymerase (<u>EP0703</u>, ThermoFisher). The oligonucleotides were purchased from Sigma-Aldrich using the default synthesis specifications (25 nmol concentration, purified by desalting). The sequences of the oligonucleotides used were: Fw 5’GCGTAATACGACTCACTATAGG ACTGGTTCTGGCAGTATTGG TTTTAGAGCTAGAAA TAGCAAGTTAAAATAAGGCTAGTC 3’ Rv 5’GATCCGCACCGACTCGGTGCCACTTTTTCAAGTTGATAACGGACTAGCCTT ATTTTAACTTGCTATTTCTAGCTCTAAAAC 3’

The amplicon was subsequently amplified using the primers: Fw: 5’ ATCCGCACCGACTCGGT 3’ and Rv: 5’ GCGTAATACGACTCACTATAG 3’ and purified using the Quick gel extraction and PCR purification combo kit (00505495, ThermoFisher). The PCR products were confirmed by an agarose gel electrophoresis and by Sanger sequencing (Base Clear, Netherlands). The sgRNA was generated using the MEGA short script ®T7 kit (AM1354, ThermoFisher) and the mRNA for a zebrafish optimized NLS-Cas9-NLS was transcribed using the mMACHINE® SP6 Transcription Kit (AM1340, Thermo Fisher) from a Cas9 plasmid (39312, Addgene) in both cases, the RNeasy Mini Elute Clean up kit (74204, QIAGEN Benelux B.V., Venlo, Netherlands) was used to purify the products. AB/TL embryos were injected with a mixture of 150 pg sgRNA /150 pg/Cas9 mRNA at 0 hpf and CRISPR injections were confirmed by PCR and Sanger sequencing. Five founders (F0) were outcrossed with AB/TL fish and efficiently transmitted the mutated allele. The chosen mutation consists of a 46 bp deletion directly after the TM1 domain and a stable line was generated by incrossing heterozygous F1 carriers. The stable homozygous *cxcr3.3* mutant line was later outcrossed with *Tg (mpeg1: mCherry-F)* and *Tg (mpx: eGFP)* transgenic lines to visualize macrophages and neutrophils, respectively.

The offspring of a *Tg (mpeg1: mCherry-F cxcr3.3+/-)* family cross was genotyped to assess the segregation pattern of the *cxcr3.3* gene. To assess macrophage and neutrophil development, a 25-30 larvae from 5 single crosses of *Tg (mpeg1: mCherry-F* WT, *cxcr3.3-/- and cxcr3.2-/-)* and *Tg (mpx: eGFP* WT, *cxcr3.3-/- and cxcr3.2-/-)* fish were pooled together and observed under a Leica M165C stereo-fluorescence microscope from 1 dpf-5 dpf to quantify the total number of macrophages and neutrophils, respectively, in the head and tail areas. The same batch of fish was observed under the stereomicroscope from 1 dpf-5 dpf to determine if there were morphological aberrations.

### Transient *cxcr3.3* overexpression

An expression construct pcDNA™3.1/V5-His TOPO-*CMV:cxcr3.3* was generated and injected into the yolk at 0 hpf to overexpress the gene in AB/TL (Figure 3-C) and *cxc3.3* mutant larvae (Figure 3-E). Overexpression levels were verified by qPCR analysis.

### Phylogenetic analysis and protein-ligand binding site prediction

Amino acid sequences of *CXCR3* genes and *ACKR*s from 13 species (Supplementary table I.) were aligned and trimmed using the free-access server gBlocks [40] and the protein evolution analysis method was fitted using ProtTest3 [41]. Evolutionary analyses were conducted in MEGA7 [42]. The evolutionary history was inferred by using the Maximum Likelihood method based on the Dayhoff matrix-based model. The tree with the highest log likelihood (−27586.19) is shown. Initial tree(s) for the heuristic search were obtained automatically by applying Neighbor-Join and BioNJ algorithms to a matrix of pairwise distances estimated using a JTT model, and then selecting the topology with superior log-likelihood value. A discrete Gamma distribution was used to model evolutionary rate differences among sites (4 categories (+*G*, parameter = 1.6611)). The tree is drawn to scale, with branch lengths measured in the number of substitutions per site. The analysis involved 48 amino acid sequences. There was a total of 529 positions in the final dataset. Protein-ligand site prediction was done using the COACH server [43, 44] and protein structure was visualized using UGENE [45, 46, 47].

### Systemic infection with *Mycobacterium marinum* and determination of bacterial burden

*M. marinum* M-strain, expressing the fluorescent marker wasabi, was grown and prepared freshly for injection as described in [48], and embryos were systemically infected with 300 CFU of *M. marinum*-wasabi by microinjection at 28 hpf in the blood island (BI) [48, 49]. Infected larvae were imaged under a Leica M165C stereo-florescence microscope and the bacterial burden was determined using a dedicated pixel counting program at 4 days post-infection (4 dpi) [50]. Data were analyzed using a two-tailed t-test and a One-way ANOVA when more than two groups were compared. Results are shown as mean ± SEM (ns p > 0.05, * p ≤ 0.05, ** p ≤ 0.01, *** p ≤ 0.001, **** p ≤ 0.0001) and combine data of 3 independent replicates of 20-30 larvae each.

### Microbicidal capacity assessment

For determining the microbicidal capacity of zebrafish larval macrophages, embryos were infected with 200 CFU of an attenuated strain, *ΔERP*-*M. marinum-*wasabi [51]. Bacteria were taken from a glycerol stock and microinjected at 28 hpf into the BI. Infected larvae were fixed with 4% paraformaldehyde PFA at 44 hpi, mounted in 1.5% low-melting-point agarose (SphaeroQ, Burgos, Spain) and bacterial clusters were quantified under a Zeiss Observer 6.5.32 laser scanning confocal microscope (Carl Zeiss, Sliedrecht, The Netherlands). A Mann-Whitney test was used to analyze the overall bacterial burden of the pooled data of 3 independent replicates of 9 fish each, where data are shown as mean ± SEM. A Kolmogorov-Smirnov test was used to analyze the distribution of bacterial cluster sizes (ns p > 0.05).

### RNA extraction, cDNA synthesis, and qPCR analysis

For every qPCR assay a total of 3 biological samples (12 larvae each) were collected in QIAzol lysis reagent (Qiagen) and RNA was extracted using the miRNeasy mini kit (Qiagen) according to manufacturer’s instructions. cDNA was generated using the iScript™ cDNA Synthesis Kit (Bio-Rad) and qPCR reactions were done using a MyiQ Single-Color Real-Time PCR Detection System (Bio-Rad) and iTaq™ Universal SYBR® Green Supermix (Bio-Rad). For every biological sample, 3 technical replicates were performed. The cycling conditions we used were: 3 min pre-denaturation at 95°C, 40 denaturation cycles for 15 sec at 95°C, annealing for 30 sec at 60°C (for all primers), and elongation for 30 sec at 72°C. All data were normalized to the housekeeping gene *ppiab* (*peptidylprolyl isomerase Ab)* and were analyzed with the 2^−ΔΔ**Ct**^ method. The following primers were used for our analyses: *ppiab* Fw: 5’ ACACTGAAACACGGAGGCAAAG 3’, *ppiab* Rv: 5’ CATCCACAACCTTCCCGAACAC 3’, *cxcr3.2* Fw: 5’ CTGGAGCTTTGTTCTCGCTGAATG 3’, *cxcr3.2* Rv: 5’ CACGATGACTAAGGAGATGATGAGCC 3’, *cxcr3.3* Fw: 5’ GCTCTCAATGCCTCTCTGGG 3’, *cxcr3.3* Rv: 5’ GACAGGTAGCAGTCCACACT 3’, *cxcl11aa* Fw: 5’ GCTCTGCTTCTTGTCAGTTTAGCTG 3’, *cxcl11aa* Rv: 5’ CCACTTCATCCATTTTACCGAGCG 3’.

A One-way ANOVA was used to test for significance and data are plotted as mean ± SEM (ns p > 0.05, * p ≤ 0.05, **p ≤ 0.01, *** p ≤ 0.001).

### Macrophage and neutrophil recruitment assays

100 CFU of *M. marinum-*wasabi (Figure 5-A-B) or 1nL of purified Cxcl11aa protein (10 ng/mL, [29]) (Figure 5-C-D) were injected into the hindbrain ventricle of *Tg (mpeg1: mCherry-F WT, cxcr3.2-/-* and *cxcr3.3-/-)* and *Tg (mpx: eGFP* WT, *cxcr3.3-/- and cxcr3.2-/-)* larvae at 48 hpf. PBS-injected larvae from each group were pooled before quantification to serve as a control group for the three genotypes. Samples were fixed with 4% PFA at 3 hpi, and macrophages within the hindbrain ventricle were counted under a Zeiss Observer 6.5.32 laser scanning confocal microscope (Carl Zeiss, Sliedrecht, The Netherlands) by going through a z-stack comprising the whole hindbrain ventricle. For the tail-amputation model, > 50 anesthetized 3 dpf larvae were put on a 2% agarose covered petri-dish and the caudal fin was cut with a glass blade avoiding to damage the notochord. Amputated larvae were put back into egg water and fixed with 4% PFA 4hours after amputation. The tail area was imaged with a Leica M165C stereo-florescence microscope and images were visualized using the LAS AF lite software. The macrophages localized within an area of 500 µm from the cut towards the trunk were counted as recruited cells (Figure 5-F). For both the hindbrain injection and the tail-amputation assays, a Kruskal-Wallis test was conducted to assess significance (* p ≤ 0.05, *** p ≤ 0.001, **** p ≤ 0.0001) and data are shown as mean ± SEM.

### Tracking of migrating macrophages

Time-lapse images of migrating macrophages from two independent replicates (5 larvae per genotype each) near the caudal hematopoietic tissue (CHT) were acquired every 2 min for 3h under basal conditions (Figure 6-A). To prevent imaging artifacts due to tail regeneration processes, time-lapse images of macrophages migrating towards the injury (3 independent replicates of 4 larvae per group each) using the tail-amputation model were acquired every 60 sec for 1.5 h (Figure 6-C). 4-5 larvae of each group and for each condition (basal/wound-induced migration) were mounted in 1.5 % low-melting-point agarose and microscopy was done using a Nikon Eclipse Ti-E microscope (Nikon Instruments Europe B.V.) with a Plan Apo 20X/0.75 NA objective. Data were saved as maximum projection images and were further analyzed using the Fiji/ImageJ [52] plugin TrackMate v3.4.2 [53]. The tracking setting used were: Log detector, estimated blob diameter=20 microns, threshold diameter= 15 microns, no further initial thresholding method was applied. The chosen view was hyperstack displayer and the tracking algorithm chosen was the simple LAP tracker, keeping the default settings. Tracks were later filtered according to the numbers of spots on track (> 40 spots / track) and spots, links, and track statistics were used to estimate the mean speed of moving macrophages and the total displacement. The total displacement was manually calculated in Excel by adding all the links of a given track and a filter was applied to classify tracks with a maximum displacement < 20microns as static cells (mean speed = 0 and total displacement = 0). Data were analyzed with a One-way ANOVA (ns p > 0.05, * p ≤ 0.05, **p ≤ 0.01) and are shown as mean ± SEM.

### Macrophage circularity assessment

The cell circularity indexes were calculated using the “analyze particle” option in the Fiji/ImageJ software [52]. The maximum projection images of migrating macrophages of the three genotypes were processed in Fiji/ImageJ by using the “despeckle” filter and by generating a binary image. In total, 30 macrophages per larvae were manually selected and the circularity of the cell in every frame was determined using the “analyze particle” option. A frequency histogram (%) for each group was plotted using cell circularity index (CI) bins as follows: 0-0.2, 0.2-0.4, 0.4-0.6, 0.6-0.8 and 0.8-1. The frequency distributions were analyzed using a Kolmogorov-Smirnov test taking the WT distribution as reference distribution (** p ≤ 0.01, **** p ≤ 0.0001).

### Bacterial dissemination assessment

200 CFU of *M.marinum-mCherry* were injected into the hindbrain ventricle of >30 WT, *cxcr3.2* and *cxcr3.3* mutants at 28 hpf. Whole larvae and tail areas were imaged with a Leica M165C stereo-fluorescence microscope and visualized with the LAS AF lite software. Images were cropped in such way that the area encompassing the tail was always the same size (4 in/10.16cm x 11 in/27.94cm). The number and size of distal granulomas were analyzed with the “analyze particle” function in Fiji/ImageJ [52]. Particles with a total area >0.002 were considered as granulomas, smaller particles were excluded from our analysis. The percentage of infected larvae that developed distal granulomas was manually calculated and a χ^2^ test was used to assess significance. A One-way ANOVA was used to assess cluster number and size (ns p > 0.05, * p ≤ 0.05, **p ≤ 0.01, ***p ≤ 0.001, ****p ≤ 0.0001). Data are shown as mean ± SEM.

### Chemical inhibition of Cxcr3.2 and Cxcr3.3

Approximately 30 3-day old larvae of each genotype (WT, *cxcr3.2-/-* and *cxcr3.3-/-)* were pre-incubated in 2 mL egg water containing either DMSO (0.01%) or NBI 74330 (50 µM) for 2 hours before tail-amputation. Larvae were put back into 2mL egg water containing either DMSO or NBI 74330 after the amputation for 4h followed by fixation with 4% PFA. Imaging of the tail region and quantification of macrophage recruitment to the tail-amputation area was done as described above. For the bacterial burden assay, approximately 30 larvae of each group were pre-incubated either with 25 µM NBI74330 or 0.01% DMSO for 3 hours before infection (24 hpf-27 hpf). Larvae were infected with 300 CFU *M. marinum-*wasabi at 28 hpf in the BI and NBI74330 and DMSO treatments were refreshed at 48 hpi. Imaging and bacterial pixel quantification were performed as described above.

## Online Supplemental Material

### Online supplementary videos 1

Representative time-lapses of WT, *cxcr3.2* mutants, and *cxcr3.3* mutant larvae after tail amputation. Time-lapses show macrophages of WT, *cxcr3.2* mutants, and *cxcr3.3* mutant larvae migrating towards the injury. Images from a z-stack of the injured area were acquired every 60 sec for 1.5 h and combined into max projection time-lapse.

## Results

### Cxcr3.3 has features of both conventional Cxcr3 receptors and ACKRs

We have previously shown that zebrafish Cxcr3.2 is a functional homolog of human CXCR3, required for macrophage migration towards the infection-inducible Cxcl11aa chemokine ligand [29]. Since macrophages also express the paralog Cxcr3.3, we set out to investigate the interaction between these two Cxcr3 family receptors. Our phylogenetic analysis revealed that Cxcr3.3 clusters in the same branch as conventional Cxcr3 chemokine receptors (Figure 1A) despite having an altered E/DRY-motif (DCY) and distinctive microswitch features of ACKRs, which are unable to conventional signaling through G-proteins (Figure 1-B). A protein-ligand binding site prediction algorithm [43, 44] showed that Cxcr3.2 and Cxcr3.3 share relevant structural features, such as a well conserved main ligand-binding site (Figure 1-C and D). While classical CXCR3 ligands (CXCL9, 10, 11) were not found, possibly due to the evolutionary distance between human and zebrafish, the top 4 hits for predicted ligands by this algorithm were shared by both Cxcr3 paralogs further confirming the well-preserved protein structure (Supplementary Table I). Since the conventional and atypical Cxcr3 paralogs cluster together, the alterations in the E/DRY-motif and in microswitches cannot be regarded as phylogenetic diagnostic features, yet these characteristics are known to be functionally determinant for GPCR activation [13, 54, 16]. Based on these observations, we hypothesize that Cxrc3.3 might antagonize the function of Cxcr3.2 since both receptors are predicted to bind the same ligands but Cxcr3.3 lacks the E/DRY-motif that is required for activation of downstream G-protein signaling and might, therefore, function as a scavenger.

**Figure 1.**
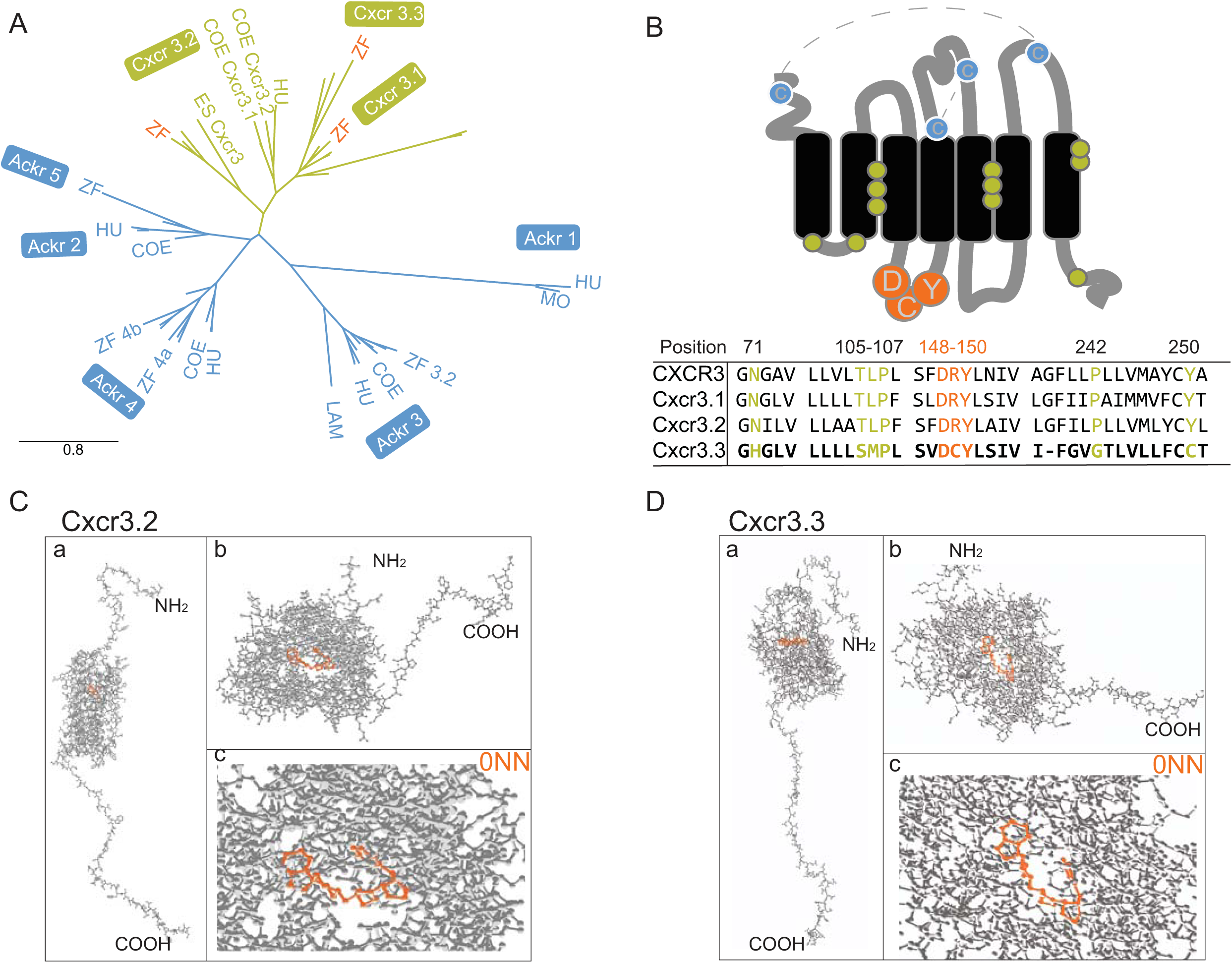
Cxcr3.3 has features of both conventional Cxcr3 receptors and ACKRs. Phylogenetic analyses including CXCR3 (green) and ACKR sequences (blue) of relevant species revealed that Cxcr3.3 is closely related to its paralogs Cxcr3.1 and Cxcr3.2 **(A)** (ZF=zebrafish, COE=coelacanth, HU=human, MO=mouse, ES= elephant shark, LAM= lamprey) despite having structural features of ACKRs **(B)**, such as an altered E/DRY-motif (orange) and microswitches (green). The predicted primary ligand-binding site of both Cxcr3.2 and Cxcr3.3 is highly conserved and structural predictions suggest that they share several ligands (Supplementary table II). **C** and **D** show the whole predicted structure of the Cxcr3.2 and Cxcr3.3 receptors **(a)**, the ligand binding site of both proteins **(b)** and the binding of one of the shared predicted ligands (0NN) by each receptor **(c)**.

### *cxcr3.3* mutant larvae do not show morphological aberrations but transient differences in macrophage development

Using CRISPR-Cas9 technology we generated a *cxcr3.3* mutant zebrafish line. The mutation consists of a 46 bp deletion in the first exon, directly after the first transmembrane domain which guarantees that the GPCR is entirely dysfunctional (Figure 2-A, B). The mutated gene did not affect survival since it segregated following Mendelian proportions (Figure 2-C). The development of mutant embryos was tracked from 24 hpf-5 dpf and no evident morphological aberrations were observed (Figure 2-D). Macrophages of *cxcr3.3* mutant and WT siblings in *Tg (mpeg1: mCherry-F)* reporter background embryos were quantified from 24 hpf-5 dpf. We also included the previously described *cxcr3.2* mutant [29] in this analysis. The total number of macrophages (Figure 2-E) in *cxcr3.3* mutant larvae was higher at day 2. However, this minor increase was short-lived since by day 3 there was no difference among the groups. We also quantified macrophages in the head and tail since these were relevant areas for our experimental setups. We observed that at day 4, *cxcr3.2*-/- larvae had transiently fewer cells in the head area (Figure 2-F). On the other hand, *cxcr3.3* mutant embryos had more macrophages during the first 2 days but leveled off after this time point (Figure 2-G). Neutrophils were quantified in the same fashion as macrophages, using a *Tg (mpx: eGFP)* reporter line, but we did not detect any difference between the groups at any time point (Supplementary figure 1.). Taking these observations into account, we performed all our experiments avoiding the time points at which macrophage development was inconsistent to prevent biased observations.

**Figure 2.**
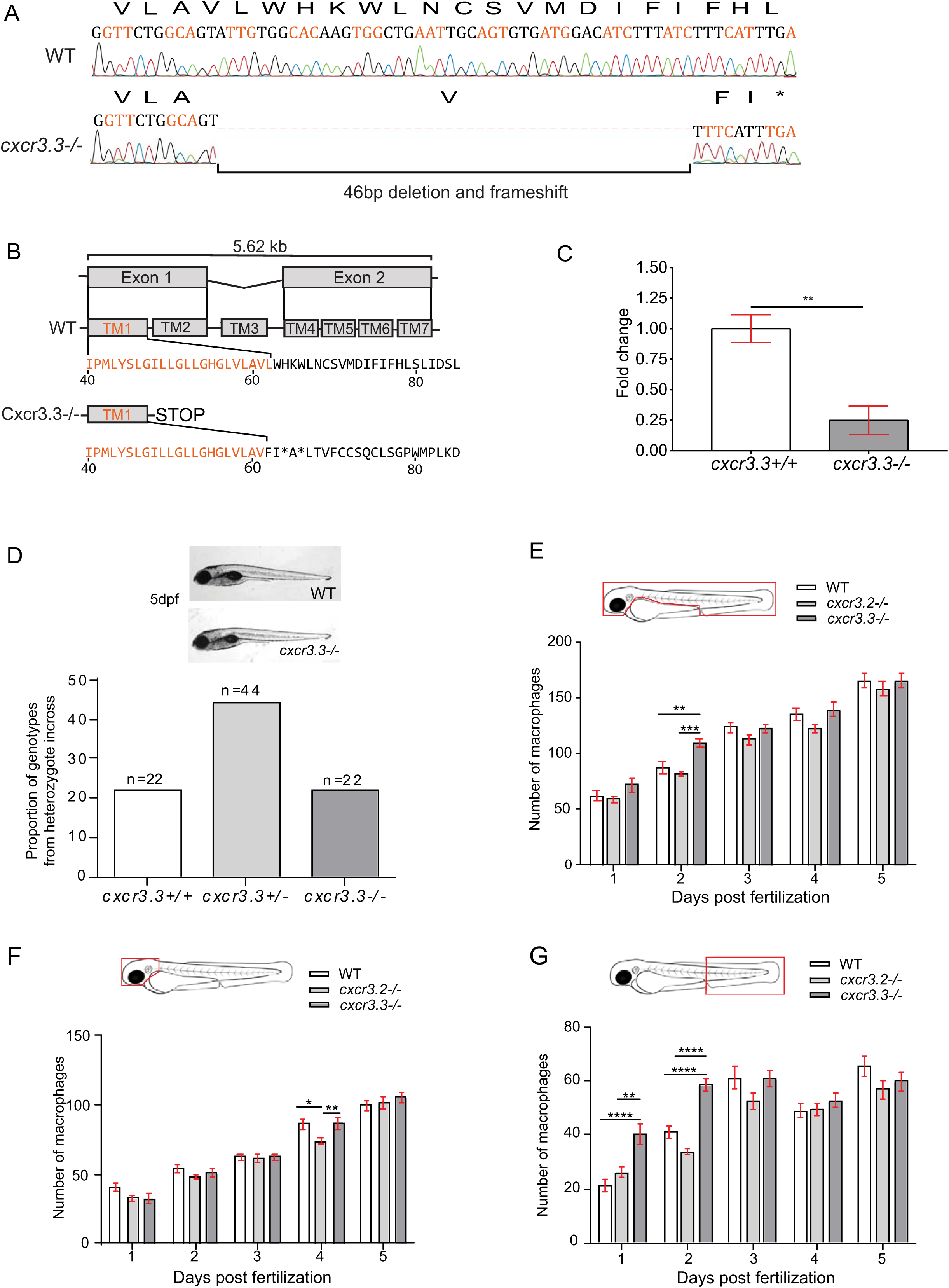
*cxcr3.3* mutant larvae do not show morphological aberrations or major differences in macrophage development. A 46 bp deletion was induced in the *cxcr3.3* gene using CRISPR-Cas9 technology **(A)**. The deletion is located in the first exon (orange), at the very end of the first transmembrane domain (TM1).The mutation shifts the reading frame and results in a premature stop codon **(B)**. Nonsense-mediated decay assessment suggests that the *cxcr3.3* mutant gene codes a truncated Cxcr3.3 protein (**C**). No evident morphological aberrations were observed in *cxcr3.3-/-* larvae within the first 5 dpf and the mutant allele segregated following Mendelian proportions **(D)**. Macrophage development was faster in *cxcr3.3-/-* embryos at 2 dpf but reverted to WT and *cxcr3.2-/-* pace after day 3 **(E)**. Fewer macrophages were found in the head area of *cxcr3.2-/-* larvae only at day 4 **(F)**, while there were more macrophages in the tail region in *cxcr3.3-/-* **(G)**. The cell numbers corresponding to each day are the average of 35 larvae of each of the 3 groups (genotypes). Data were analyzed using a two-way ANOVA and are shown as mean ± SEM (ns p > 0.05, * p ≤ 0.05, **p ≤ 0.01, *** p ≤ 0.001, **** p ≤ 0.0001).

### Deficiency of *cxcr3.3* results in a higher *M. marinum* infection burden while overexpressing the gene lowers bacterial burden

We previously showed that mutation of *cxcr3.2* enabled zebrafish larvae to better control *M. marinum* infection, a phenotype that could be explained by a reduction of macrophage migration in the absence of Cxcr3.2, which limits the dissemination of infection [29]. To investigate our hypothesis that Cxcr3.2 and Cxcr3.3 might have opposing functions, we started by determining if Cxcr3.3 was also involved in the immune response to *M. marinum*. In contrast to the effect of the *cxcr3.2* mutation, systemically infected *cxcr3.3* mutant larvae had a higher bacterial burden than WT 4 days after infection with *M. marinum* (Figure 3-A, B). We transiently overexpressed *cxcr3.3* by injecting a *CMV*: *cxcr3.3* construct into AB/TL fish at 0 hpf and observed that larvae overexpressing *cxcr3.3* had a lower bacterial burden than non-injected controls (Figure 3-C, D).To rescue the mutant phenotype, we injected the CMV: *cxcr3.3* construct into *cxcr3.3* mutant larvae. We observed that the bacterial burden of the rescued mutants (*cxcr3.3 mutants* + CMV-*cxcr3.3*) was similar to WT and significantly lower than in non-injected *cxcr3.3* mutants (Figure 3-E, F). For this assay, we used non-injected larvae as controls since there was no significant difference in bacterial burden of larvae injected with the empty CMV:vector and non-injected larvae (Supplementary figure 2.). To assess whether there was genetic compensation when one of the *cxcr3* paralogs was depleted, we assessed the gene expression of *cxcr3.2* in *cxcr3.3* mutants and *cxcr3.3* in *cxcr3.2* mutants under basal conditions and upon infection with *M. marinum*. The expression of *cxcr3.2* remained unaffected in the absence of *cxcr3.3* and was induced upon infection with *M. marinum* (Figure 3-G). On the other hand, *cxcr3.3* expression was lower in *cxcr3.2* mutant larvae and it was moderately induced upon infection (Figure 3-H). We also assessed the expression of the Cxcl11aa ligand, as it is the most upregulated one out of the 7 Cxcl11-like ligands during *M. marinum* infection, in both *cxcr3* mutants [29, 31]. The gene was induced upon infection independently of the expression on *cxcr3.2* and *cxcr3.3* (Figure 3-I). Thus, the expression of *cxcr3.3* is partially dependent on *cxcr3.2*, but it is not strongly induced upon infection like *cxcr3.2* and *cxcl11aa.* Furthermore, the expression data indicate that the increased bacterial burden of *cxcr3.3* mutants is not due to altered *cxcr3.2* expression. Together with our previous study on *cxcr3.2* [29], we conclude that mutation of *cxcr3.2* and *cxcr3.3* results in opposite infection outcomes and that *cxcr3.3* overexpression phenocopies the host-protective effect of the *cxcr3.2* mutation.

**Figure 3.**
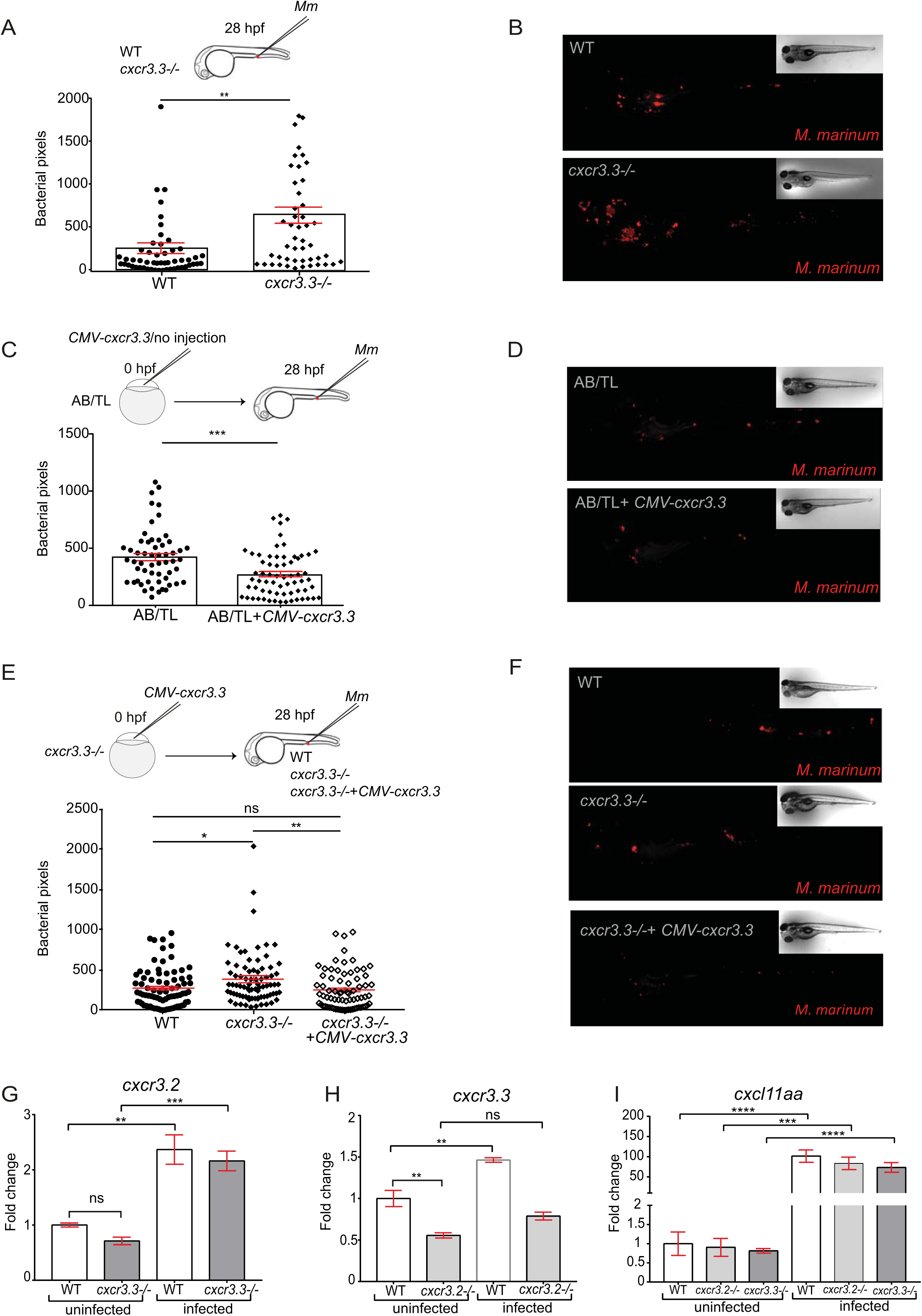
Depletion and overexpression of *cxcr3.3* result in opposite *M. marinum* infection outcomes. Cxcr3.3 deficient larvae had a higher bacterial burden than their WT siblings at 4 days following blood island (BI) infection with 300 CFU of *M. marinum (Mm)* **(A, B)**. We transiently overexpressed *cxcr3.3* in AB/TL embryos by injection of a *CMV: cxcr3.3* construct at 0 hpf and observed that bacterial burden was lower in larvae overexpressing the gene than in non-injected controls at 4 dpi **(C, D)**. To rescue the *cxcr3.3-/-* phenotype we restored the expression of the gene by transiently overexpressing it (*CMV: cxcr3.3*) in one half of the *cxcr3.3* mutants (*cxcr3.3-/-* rescued). The bacterial burden was lower in the rescued group than in non-injected *cxcr3.3* mutants (*cxcr3.3-/-*) and similar to the bacterial burden in WT controls **(E, F)**. Results from qPCR show that *cxcr3.2* expression remained unaltered in the *cxcr3.3* mutants and that it was induced upon infection **(G)**, while *cxcr3.3* expression was lower in *cxcr3.2-/-* and was moderately induced upon *Mm* infection **(H)**. The ligand *cxcl11aa* was induced upon infection independently of any of the *cxcr3* genes. In all cases, systemic infection was done at 28 hpf in the BI with 300 CFU of *Mm*. The bacterial burden data were analyzed using a two-tailed t-test **(A-C)** and a One-way ANOVA **(E).** Results are shown as mean ± SEM (ns p > 0.05, * p ≤ 0.05, **p ≤ 0.01, *** p ≤ 0.001, **** p ≤ 0.0001) and combine data of 3 independent replicates of 20-30 larvae each. The qPCR data were analyzed with the 2^−ΔΔ**Ct**^ method and a One-way ANOVA. Results are plotted as mean ± SEM (ns p > 0.05, * p ≤ 0.05, ** p ≤ 0.01, *** p ≤ 0.001).

### Macrophages lacking Cxcr3.3 efficiently clear intracellular bacteria

Lysosomal degradation of intracellular bacteria by macrophages is crucial for the containment of mycobacterial infections. The *ERP* (exported repetitive protein) virulence factor is required for bacteria to survive and replicate inside acidic compartments. Mycobacteria lacking *ERP* are easily eliminated by macrophages and can be used as an indicator of bacterial clearance efficiency since the initial infection dose (200 CFU) remains unchanged in the absence of bacterial replication [51]. To evaluate if the poor contention of the infection in *cxcr3.3 mutants* was associated to a deficient clearance of bacteria, we injected *ΔERP M. marinum* into the circulation of WT and mutant larvae and quantified bacterial clusters in the tail area at 2 dpi. Figure 4-A shows no difference between WT and mutants regarding the total number of bacterial clusters in the tail area. We divided bacterial clusters into three groups according to the number of bacteria they contained: 1-5 bacteria (small cluster), 6-10 bacteria (medium-sized cluster) and > 10 (large cluster) as shown in the representative image illustrating the cluster size categories in Figure 4-B. The frequency distributions of the 3 different cluster sizes in each genotype were compared and no significant difference was found (Figure 4-C). Mycobacterial clearance remained unaffected in the absence of Cxcr3.3, suggesting that the poor control of the infection in *cxcr3.3* mutants is not due to a deficient bacterial clearance. As a positive control, we also ran this assay using DNA-damage regulated autophagy modulator 1 *(dram1)* mutant larvae, and their WT siblings, since *dram1* mutants cannot efficiently clear *Mm* [37]. The total number of clusters was higher in *dram1* mutants and large bacterial clusters were more frequent (Supplementary figure 3.). Therefore, we conclude that macrophages in *cxcr3.3* mutants, contrary to *dram1* mutants, are not affected in their microbicidal capacity.

**Figure 4.**
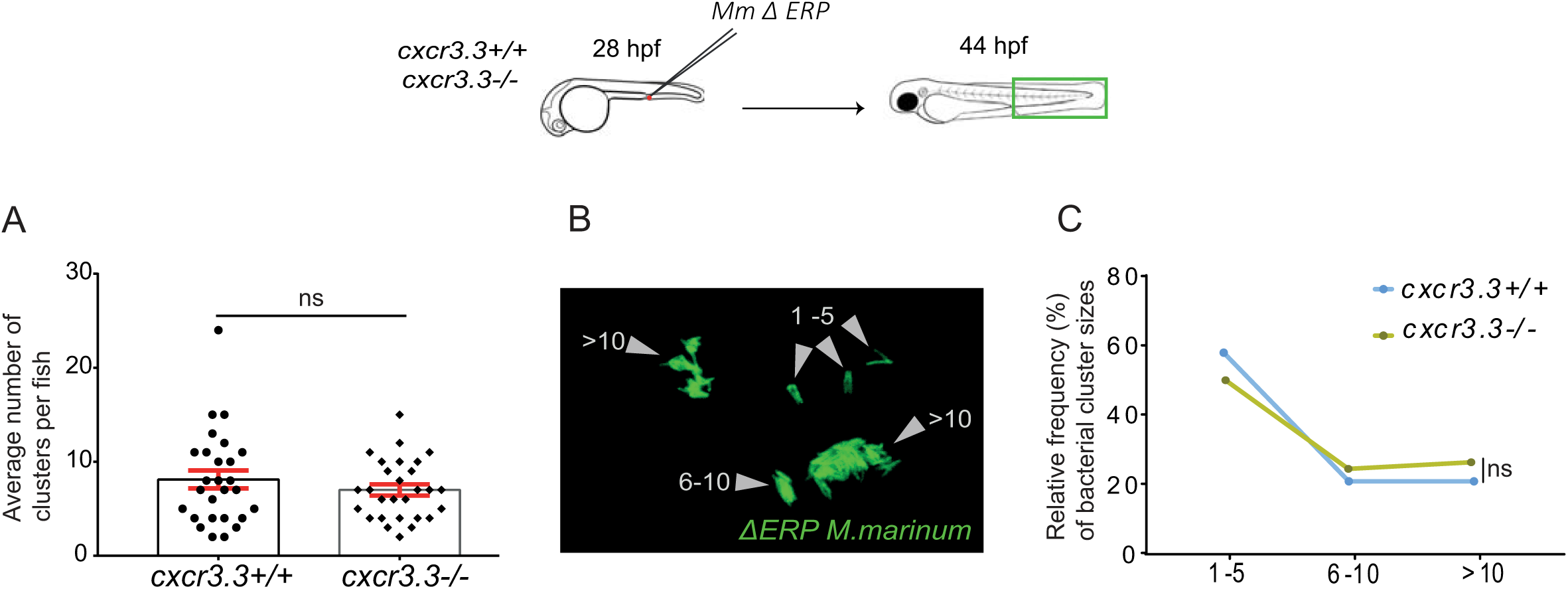
Macrophages lacking Cxcr3.3 efficiently clear intracellular bacteria. Cxcr3.3 deficient larvae and their WT siblings were infected in the BI at 28 hpf with 200 CFU of the *ΔERP M. marinum-*wasabi strain that is unable to survive and replicate inside acidic compartments and can be easily cleared by macrophages. The total number of bacterial clusters in every fish was quantified **(A)**. We divided the bacterial clusters into 3 groups based on the number of bacteria they contained (1-5,1-6 and > 10) to assess bacterial clearance at 44 hpi **(B)**. No difference between WT and *cxcr3.3-/-* cluster size distributions (frequency in %) was found **(C)**. A Mann-Whitney test was conducted to analyze the overall bacterial burden of the pooled data of 3 independent replicates of 9 fish each. Data are shown as mean ± SEM **(A)**. A Kolmogorov-Smirnov test was used to analyze the distribution of bacterial cluster sizes **(C)** (ns p > 0.05).

### Cxcr3.3-deficient macrophages show enhanced recruitment to sites of infection, towards Cxcl11aa, and to sites of injury

Several studies have shown that macrophage recruitment is essential for bacterial clearance and containment during mycobacterial pathogenesis but supports bacterial dissemination and granuloma formation at early stages of the infection [55, 56]. We previously found that *cxcr3.2* mutant larvae showed an attenuated recruitment of macrophages to sites of infection and towards Cxcl11aa ligand. This study suggested that macrophage-mediated dissemination of bacteria was reduced due to this recruitment deficiency in *cxcr3.2* mutants since fewer cells would become infected with *M. marinum* [29]. We addressed cell recruitment to examine whether the process was altered in absence of the Cxcr3.3 receptor. We infected 2-day old larvae in the hindbrain ventricle with either *M. marinum* or Cxcl11aa protein and quantified the macrophages that infiltrated into the cavity at 3 hpi. In both cases, we observed an enhanced recruitment to the site of injection in *cxcr3.3* mutants (Figure 5-A-D). In contrast, recruitment was attenuated in *cxcr3.2* mutants (Figure 5-A-D), in line with our previous results [29]. The response to mechanical damage was also assessed using the tail-amputation model. The tail fins of WT, *cxcr3.2* mutant and *cxcr3.3* mutant larvae were dissected and macrophages within an area of 500 µm from the cut towards the trunk were quantified as recruited cells. Here too, we observed opposing results between the Cxcr3 mutants: more cells were recruited in the *cxcr3.3* mutants and fewer cells were recruited to the site of damage in the *cxcr3.2* deficient larvae (Figure 5-E, F). We conclude that Cxcr3.3 and Cxcr3.2 deficiencies have opposing phenotypes regarding macrophage recruitment to sites of infection and injury or to a source of Cxcl11aa chemokine. While attenuated macrophage recruitment in *cxcr3.2* mutants favors bacterial contention [29], the enhanced recruitment of macrophages to sites of infection in *cxcr3.3* mutants might be facilitating macrophage-mediated dissemination of bacteria, resulting in the increased bacterial burden observed in our infection experiments.

**Figure 5.**
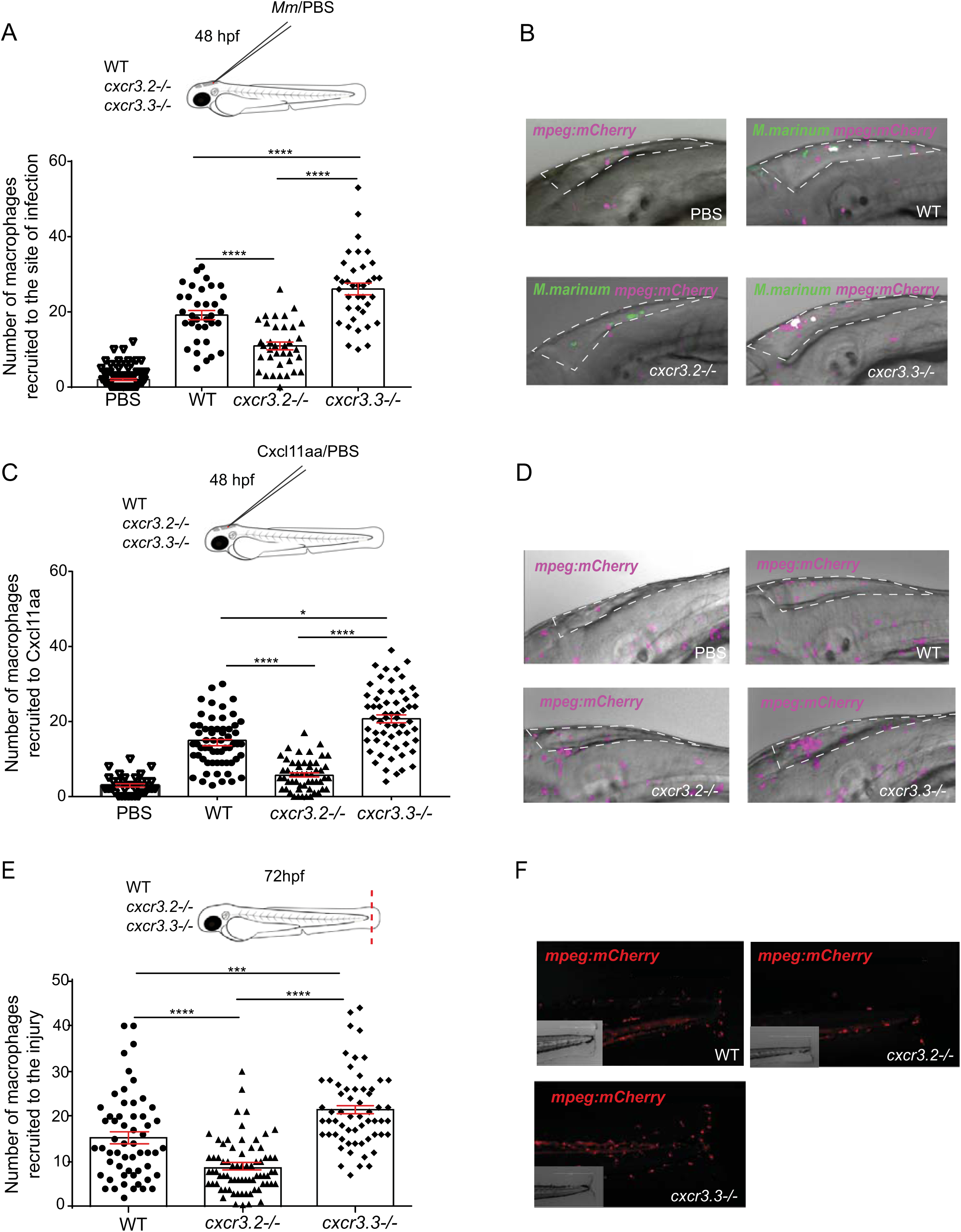
Macrophages lacking Cxcr3.3 show an enhanced recruitment to sites of infection, towards Cxcl11aa, and to sites of mechanical damage. Significantly fewer cells were recruited to the hindbrain ventricle in *cxcr3.2-/-* at 3 hpi with *Mm* and more macrophages were recruited to the same site in *cxcr3.3-/-* compared to WT controls **(A, B)**. The same trend was observed when 1nL of Cxcl11aa protein (10 ng/mL) was injected in the same experimental setup **(C, D)**. To assess macrophage recruitment to sites of injury, we used the tail-amputation model and observed enhanced recruitment of macrophages in *cxcr3.3-/-* larvae and attenuated recruitment of macrophages in *cxcr3.2-/-* relative to WT at 4 hpa **(E, F)**. The PBS injected control group combines WT, *cxcr3.2* and *cxcr3.3* mutants and shows no cell recruitment at 3 hpi. In all cases, statistical analyses were done with pooled data of three independent replicates (20-30 larvae per group each). A Kruskal-Wallis test was used to assess significance (* p ≤ 0.05, *** p ≤ 0.001, **** p ≤ 0.0001) and data are shown as mean ± SEM.

### Cxcr3.3 depletion has no significant effect on neutrophil recruitment to sites of infection or injury

Although macrophages are the first responders towards mycobacterial infection and the main components of granulomas, neutrophils are also recruited to infectious foci and participate in the early immune response [57, 58]. Besides, both Cxcr3.2 and Cxcr3.3 are also expressed on this cell-type [29]. Therefore, we assessed the effect of the *cxcr3.2* and *cxcr3.3* mutations on neutrophil recruitment to local *Mm* infection and upon injury similar as for macrophages in the previous section (Figure 6). When *Mm* was locally injected into the hindbrain, fewer neutrophils were recruited to the cavity in *cxcr3.2* mutants at 3 hpi, while there was no difference between WT and *cxcr3.3* mutants (Figure 6 A-B). Using the tail-amputation model to assess cell recruitment, we observed the same pattern: the lack of *cxcr3.2* reduced neutrophil recruitment to injury while recruitment remained unaffected in *cxcr3.3* mutants (Figure 6 C-D). Our data suggest that Cxcr3.2 is required for neutrophil recruitment, as shown by previous studies [59], and that the effect of the *cxcr3.3* mutation does not significantly impact the migratory properties of this cell type.

**Figure 6.**
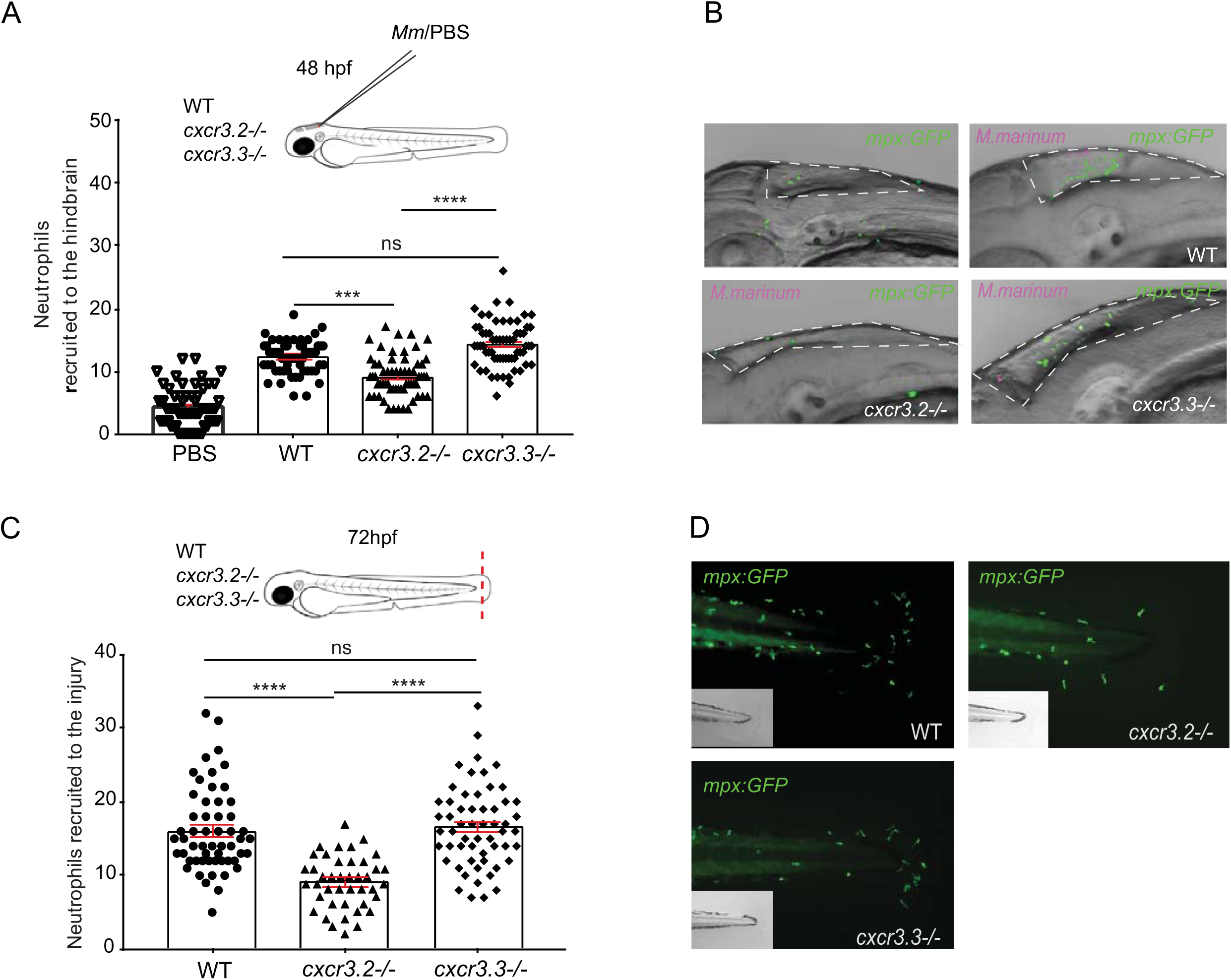
Neutrophil recruitment to sites of infection and injury is not altered in *cxcr3.3* mutants. 100 CFU of *Mm*-mCherry were injected in the hb ventricle of 2 day-old WT, *cxcr3.2* and *cxcr3.3* mutant larvae to assess neutrophil (*mpx: eGFP*) recruitment to the infection site at 3 hpi. The number of cells that infiltrated the cavity was lower in *cxcr3.2* mutants but remained unchanged in WT and *cxcr3.3* mutants (A-B). The tail fin of WT larvae and *cxcr3.2* and *cxcr3.3* mutants was amputated and neutrophil recruitment was assessed at 4 hours post amputation. There were fewer recruited neutrophils in the *cxcr3.2* mutants while there was no difference between *cxcr3.3* mutants and WT. The PBS injected control group PBS combines WT, *cxcr3.2* and *cxcr3.3* mutants and shows no cell recruitment at 3 hpi. In all cases, statistical analyses were done with pooled data of three independent replicates (20-30 larvae per group each). A Kruskal-Wallis test was used to assess significance (ns p > 0.05, *** p ≤ 0.001, **** p ≤ 0.0001) and data are shown as mean ± SEM.

### Macrophages lacking Cxcr3.3 move faster than WT cells under basal conditions and upon mechanical damage, and have an elongated and branched morphology

We previously reported that macrophage recruitment to sites infection was attenuated in *cxcr3.2* mutant macrophages because cells were less motile [29]. To further examine the role of cell recruitment in *M. marinum* pathogenesis, we assessed if macrophage motility was also affected when Cxcr3.3 was ablated. Cell motility was reviewed concerning total cell displacement and average speed. No significant difference was found in total cell displacement under basal conditions (Figure 7-A, B-1) but *cxcr3.3* mutant macrophages moved faster than the other two groups (Figure 7-A, B-2). To induce directional migration of macrophages we used the tail-amputation model. The tracks covered by *cxcr3.2* mutant macrophages were shorter when we induced directed migration (Figure 7-C, D-1). In contrast, Cxcr3.3-deficient macrophages moved faster than the remaining groups when the tail-amputation model was employed (Figure 7-C, D-2, Supplementary videos 1.). Cell circularity index (CI) was assessed as an indicator of motility and activation status of macrophages. Both *cxcr3* mutant CI distributions differ from the WT. The distribution of the CI values of Cxcr3.3-depleted macrophages shows that more cells are branched and elongated, while the CI value distribution in the *cxcr3.2* mutants suggests that macrophages are rounder (Figure 6-E). The most frequent CI interval within WT macrophages was 0.4-0.6 (42%), for *cxcr3.2* mutants it was 0.4-0.8 (71%) and for *cxcr3.3* mutants, 0.2-0.4 (39%) (Figure 7-F). To further confirm that *cxcr3.2* and *cxcr3.3* mutants have different activation profiles we assessed the transcriptional profile of the inflammatory markers *tnfa*, *cxcl11aa* and *il1b* at 4 hours post amputation in the three groups and found that *tnfa* and *cxcl11aa* were upregulated in *cxcr3.3* mutants (Supplementary figure 4.). Taken together, these data suggest that macrophage recruitment in *cxcr3.3* mutants results from a faster displacement of these cells to reach sites of infection or other inflammatory stimuli. This increased speed is linked to a higher macrophage activation status (lower CI values) and a pro-inflammatory phenotype of the *cxcr3.3* mutant fish. Therefore, we propose that the progression of *M. marinum* infection is accelerated in *cxcr3.3* mutants by facilitating the spreading of bacteria into newly recruited macrophages and the subsequent seeding of secondary granulomas

**Figure 7.**
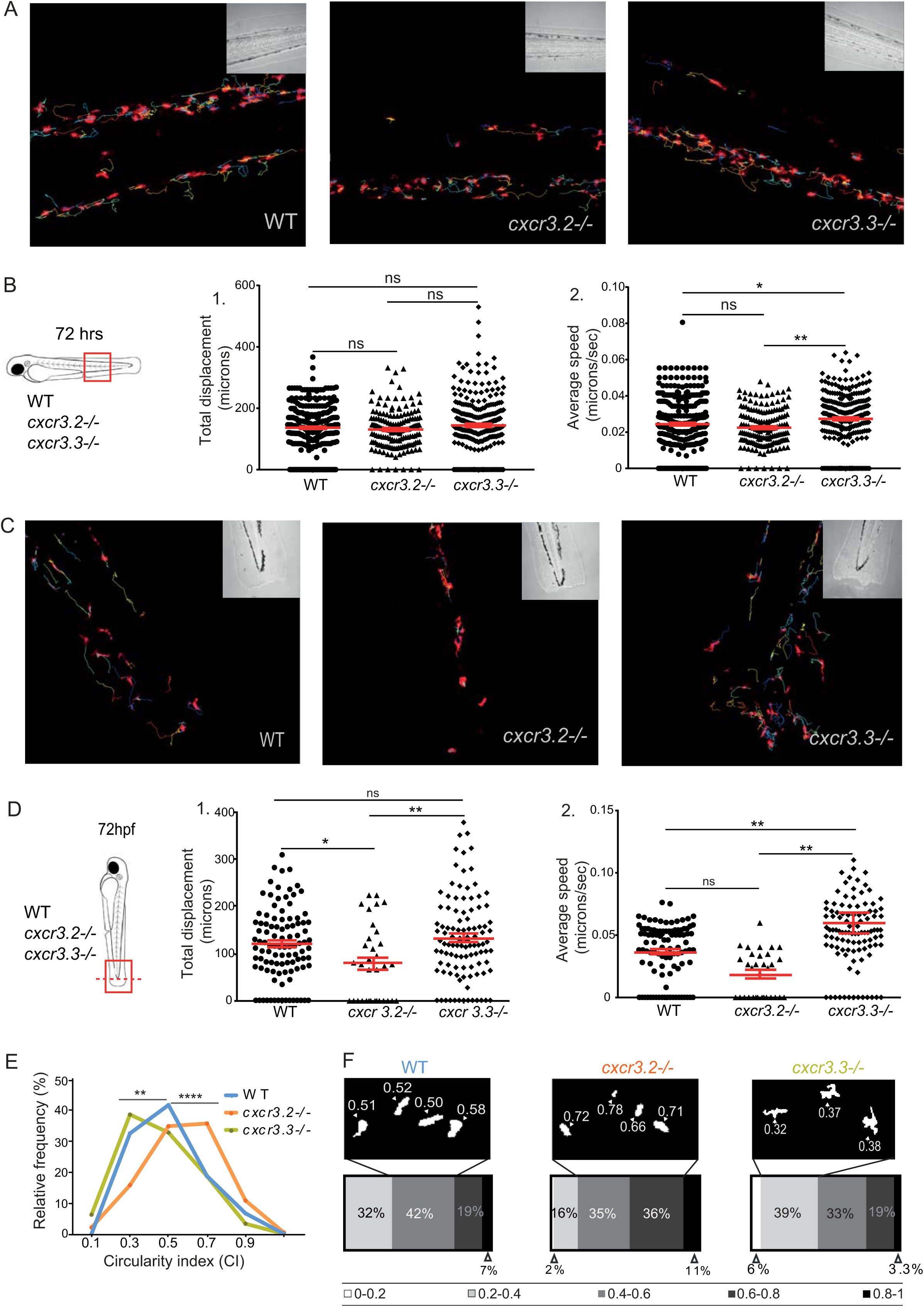
Cxcr3.3 depleted macrophages move faster than WT cells under basal conditions and upon mechanical damage and have a lower circularity index (CI). Panel **A** shows representative images of tracks of macrophages of 3-day-old larvae from the three genotypes under unchallenged conditions (random patrolling). Macrophages were tracked for 3 h and images were taken every 2 minutes. Graphs in **B** show the total displacement of all cells tracked shortly after amputation in each group throughout 3h **(B-2)** and the average speed of each cell **(B-2)**. In this case, macrophages were tracked for 1.5 h and images were acquired every 1 minute. There was no significant difference between the groups in terms of total cell displacement (**B-1.**), however *cxcr3.3-/-* macrophages did move faster than the remaining groups as indicated by the dot-plots in (**B-2.**). Panel **C** shows representative images of macrophage tracks of the three groups directly after a tail amputation. The tracks of *cxcr3.2-/-* macrophages were shorter than those of the remaining groups **(D-1.)** and *cxcr3.3-/-* macrophages moved faster than the other two groups when mechanical damage was inflicted **(D-2.)**. Data of unchallenged larvae were collected from two independent replicates (5 larvae per group each) and for the tail-amputation model data from 3 independent replicates (4 larvae per group each) were pooled together for analysis. A One-way ANOVA was performed to test for significance (ns p > 0.05, * p ≤ 0.05, ** p ≤ 0.01) and data are shown as mean ± SEM. The circularity index (CI) distributions of both *cxcr3.2-/-* and *cxc3.3-/-* differ from the WT control but are skewed in opposite directions as low CI values are more frequent in *cxcr3.3* mutants than in WT and high CI values are more frequent in *cxcr3.2* mutants as shown by the curves. **(E)**. Panel **F** shows representative images of the most frequent CI interval in each group and the bar displays the percentage of each CI category within each genotype. A Kolmogorov-Smirnov test was used to evaluate the CI value distributions using the WT data as reference distribution (** p ≤ 0.01, **** p ≤ 0.0001).

### Enhanced motility of *cxcr3.3* mutant macrophages facilitates cell-mediated *M. marinum* dissemination

Taking into account that *cxcr3.3* mutant macrophages move faster and are recruited more efficiently to sites of infection, we wanted to know whether the enhanced motility of macrophages in *cxcr3.3* mutants could facilitate bacterial dissemination by accelerating granuloma formation and seeding of secondary granulomas. We addressed our question by locally injecting *Mm* into the hindbrain of WT, *cxcr3.2* and *cxcr3.3* mutants at 28 hpf and by assessing the percentage of infected larvae that developed distal granulomas at 4 dpi (Figure 8-A), as well as the number and size of such granulomas in each group (Figure 8-C,D). Our data show that *cxcr3.3* mutant larvae more frequently developed distal granulomas (22%) than the other two groups (Figure 8-A). In addition, the average number of the distal granulomas per fish within this group was higher (Figure 8 C) and the quantified structures were also larger (Figure 8-D). Consistent with previous work [29], a small proportion of *cxcr3.2* mutant larvae (5%) developed fewer and smaller distal granulomas compared with the wild type (12.7%). Our data suggest that *cxcr3.3* mutant macrophages favor bacterial dissemination and the seeding of secondary granulomas due to their enhanced recruitment to sites of infection and their higher speed.

**Figure 8.**
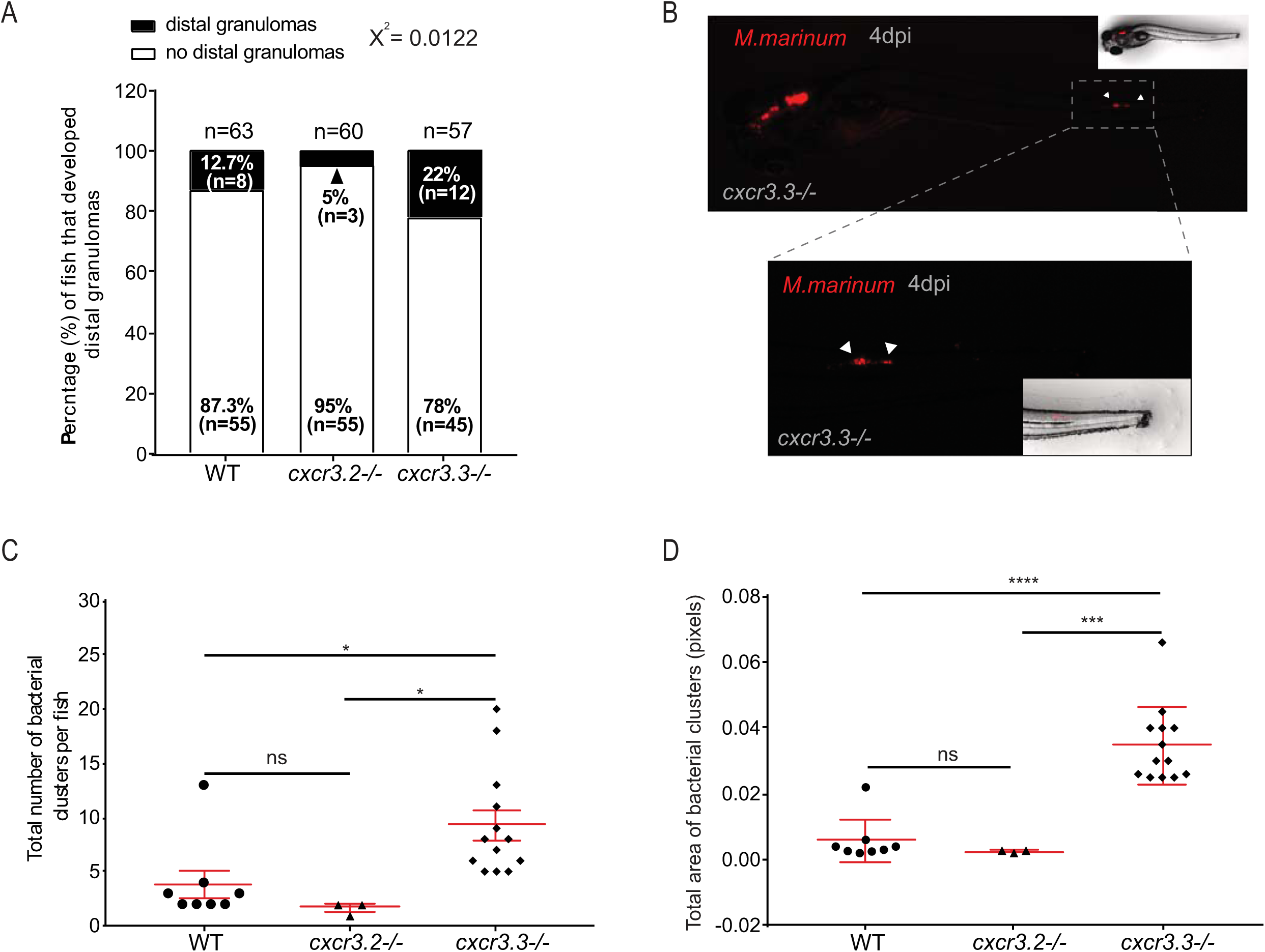
Enhanced motility of *cxcr3.3* mutant macrophages facilitates bacterial dissemination. Four days after local infection with 200 CFU of *Mm* in the hb, *cxcr3.3* mutants developed more distal granulomas (22%) than WT (12.7%) and *cxcr3.2* mutants (5%) while the latter developed fewer than the other two groups (A). Embryos from the three genotypes were infected at 28 hpf and imaged under the stereo fluorescence microscope (whole body and zoom to the tail) at 4 dpi. **B** illustrates the imaging process of a representative *cxcr3.3* mutant larvae. Cxcr3.3 depleted larvae developed more distal granulomas per fish (**C**) and these granulomas were also larger in *cxcr3.3* mutants than the other two groups, while *cxcr3.2* mutants showed an opposite trend (**D**). Graphs show pooled data from four independent replicates, each of 12-15 infected larvae per group. The number and size of distal granulomas were determined using the “analyze particle” function in Fiji. A χ^2^ test was conducted to assess differences in the proportion of larvae that developed distal granulomas within the 3 groups and a One-way ANOVA to compare the number and size of distal granulomas (ns p > 0.05, * p ≤ 0.05, *** p ≤ 0.001 and **** p ≤ 0.001). Data are shown as mean ± SEM.

### Chemical inhibition of both Cxcr3 receptors affects only macrophages expressing Cxcr3.2 and phenocopies *cxcr3.2* mutants regarding bacterial burden and macrophage recruitment efficiency

To further inquire into the roles and interactions of Cxcr3.2 and Cxcr3.3 we chemically inhibited both receptors simultaneously and addressed macrophage recruitment using the tail-amputation model and the *M. marinum* infection model. To this end, we used the allosteric CXCR3-specific inhibitor NBI 74330, of which the binding site is highly conserved in the Cxcr3.2 and Cxcr3.3 protein structures [29]. WT, *cxcr3.2* mutant and *cxcr3.3* mutant larvae were bath-exposed for 3 h before amputation and for another 4 h following tail-amputation to NBI 74330 (50 µM) or to the vehicle DMSO (0.05%) as a control. A significant reduction in the number of recruited macrophages occurred in WT and *cxcr3.3* mutants, but there was no decline in cell recruitment in *cxcr3.2* mutants when exposed to the inhibitor (Figure 9 A-D). Similarly, WT larvae were bath exposed to NBI 74330 (25 µM) for 3h prior systemic infection with *M. marinum* and kept in NBI 74330 (25 µM) for the following 4 days. Inhibition of both Cxcr3 receptors resulted in a lower bacterial burden than that of larvae treated with DMSO (Figure 9-E-F) and thereby phenocopied the effects of the *cxcr3.2* mutation [29] or *cxcr3.3* overexpression (this study). These results support our hypothesis that Cxcr3.2, an active GPCR, is essential for macrophage motility and recruitment to different stimuli while Cxcr3.3, an ACKR with predicted scavenger function, does not play a direct role on these processes but indirectly regulates them by competing with active receptors for shared ligands.

**Figure 9.**
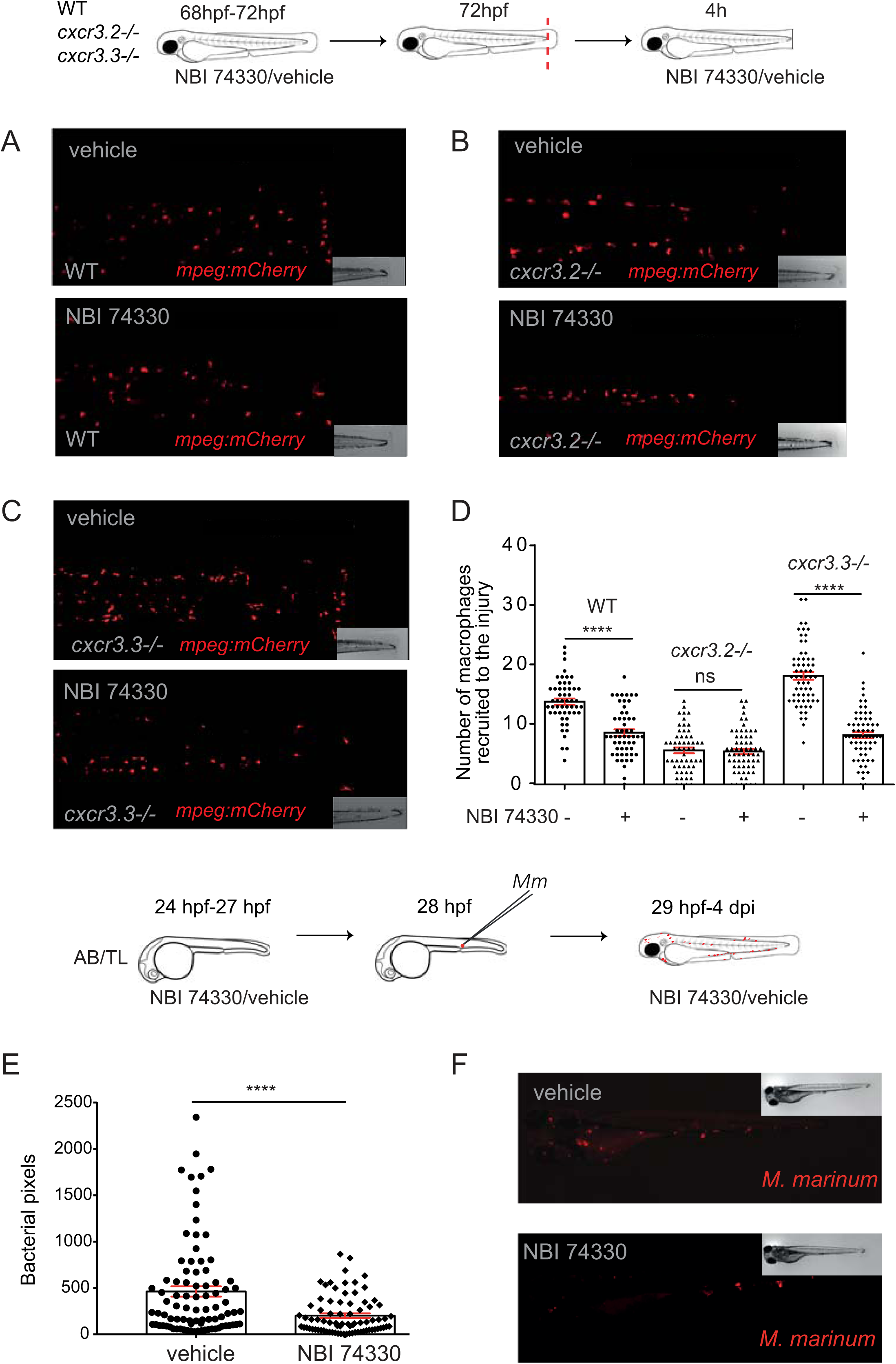
Chemical inhibition of both Cxcr3 receptors affects only macrophages expressing Cxcr3.2 and renders a similar bacterial burden and macrophage recruitment efficiency as *cxcr3.2* mutants. After bath exposure of 3-day old larvae to either the CXCR3-specific inhibitor NBI 74330 (50 µM) or vehicle (DMSO 0.01%), before and after tail-amputation showed that cell recruitment to the site of injury was reduced in macrophages expressing Cxcr3.2, namely WT and *cxcr3.3-/-* **(A, C)**, while no further decline in cell recruitment was observed in *cxcr3.2* mutants **(B, D)**. Chemical inhibition of both Cxcr3 receptors with NBI 74330 (25 µM) before and after systemic infection with *Mm* resulted in a lower bacterial burden at 4 dpi than in the vehicle-treated control and resembles the *cxcr3.2* mutant phenotype **(E, F)**. The data of three independent replicates were pooled and are presented as mean ± SEM. A Kruskal-Wallis test was conducted to assess significance (ns p > 0.05, **** p ≤ 0.0001) in the macrophage recruitment assay. Only the p values between each condition (vehicle/ NBI 74330) within each group are shown **(D)**. Bacterial burden data were analyzed using a two-tailed t-test and data are shown as mean ± SEM (**** p ≤ 0.0001).

## Discussion

Our findings illustrate the evolution of regulatory mechanisms in chemokine signaling networks and show how positive or negative dysregulation of the CXCR3 signaling axis results in opposite outcomes on macrophage behavior and innate host defense against mycobacterial infection. The *Mycobacterium tuberculosis* epidemiology highlights the urgent need to develop new clinical strategies to treat the infection given the growing incidence of multidrug-resistant strains and the high prevalence of the infection worldwide [12, 60]. GPCRs, such as chemokine receptors, are the largest protein family targeted by approved pharmacological therapies [61]. Therefore, it is important to further our understanding of the fundamental regulatory mechanisms of GPCR-related pathways. In the present study, we used the zebrafish model to functionally characterize the antagonistic interplay between two CXCR3 paralogs in the context of mycobacterial infection and mechanical damage. Our results suggest that the potential scavenging activity of an atypical CXCR3 paralog, Cxcr3.3, fine-tunes the activity of the functional CXCR3 paralog, Cxcr3.2, serving as a regulatory mechanism for the modulation of the immune response. These findings highlight the relevance of ACKRs as regulatory components of chemokine signaling networks.

At present, 5 ACKRs have been described in vertebrates, namely, ACKR1 (DARK), ACKR2, ACKR3 (CXCR7), ACKR4, and ACKR5 [12, 18]. The identification of ACKRs and the subsequent classification of these receptors within the subfamily is complex given their structural heterogeneity and the limited phylogenetic homology [17, 18, 15]. However, as in all GPCRs, the E/DRY motif and microswitch elements are indicative of the activation status and function of a receptor [16]. Microswitches stabilize the active conformation of GPCRs and are highly conserved residues, which are unchanged in Cxcr3.2 but not in Cxcr3.3 [13, 16]. The Asp (D) and the Arg (R) of the E/DRY-motif are key residues to stabilize the inactive conformation of GPCRs by forming a salt bridge between the 3^rd^ IC loop and TM6 that blocks G-protein coupling. This so-called “iconic-lock” breaks upon binding of an agonist and triggers structural rearrangements that expose the G-protein docking site and enables canonical (G-protein-dependent) downstream signaling [16]. Substitutions in the E/D and Y within the E/DRY-motif are commonly associated with the permanent activation of the receptor and gain of function events, while substitutions of the R, as found in Cxcr3.3 (DCY motif), have been shown to result in the permanent “locking” of the receptor and a consequent loss of function [16, 54, 62]. The E/DRY motif also interacts with the intracellular COOH terminus that is critical for GPCR activation and with Gα subunits. It is noteworthy to mention that chemokine receptors can also signal in a G-protein-independent manner through β-arrestin in the context of chemotaxis, and that this pathway is associated with the internalization and subsequent intracellular degradation of ligands [16, 62]. Altogether, this information led us to propose that Cxcr3.3 is an ACKR The zebrafish genome encodes a family of seven *cxcl11*-like paralogous genes, which are thought to share common ancestry with *CXCL9-10-11*, the ligands of human CXCR3 [29]. We have previously shown that one of the *cxcl11*-like genes, *cxcl11aa*, is strongly inducible by mycobacterial infection and by mechanical damage [29, 63]. Subsequently, we used an *in vivo* macrophage migration assay in *cxcr3.2* mutants and wild-type siblings to demonstrate that purified Cxcl11aa protein acts as a ligand for the Cxcr3.2 receptor. Based on the structural conservation of the ligand binding site Cxcr3.3 is predicted to bind the same ligands as Cxcr3.2. This is consistent with several studies reporting that mutations in GPCRs may affect the structure of the receptor preventing the opening of the intercellular cavity required for G-protein coupling and subsequent signaling, while ligand affinity remains unchanged [16].

Furthermore, the fact that the top hits in our ligand prediction analysis are shared by both Cxcr3 paralogs strongly suggests that both receptors can bind the same ligands due to the highly conserved hydrophobic residues in the ligand-binding site. Studies showing that signaling by CXCL11 and CXCL12 chemokines is subject to ACKR regulation [13, 18], set a precedent for our hypothesis that both receptors can bind the same ligands but only Cxcr3.2 can signal in a canonical manner, while Cxcr3.3 acts as a regulator by scavenging shared ligands. Nevertheless, biochemical data are required to fully confirm our hypothesis.

To deconstruct the proposed antagonism of Cxcr3.3 on Cxcr3.2 activity, we first compared the overall outcomes of *M. marinum* infection in *cxcr3.2* and *cxcr3.3* mutants. In agreement with our hypothesis, we observed that *cxcr3.2* and *cxcr3.3* mutants have opposite infection phenotypes. While our previous results showed that *cxcr3.2* mutants have increased resistance to mycobacterial infection [29], similar to *cxcr3* mutant mice [30], the *cxcr3.3* mutant generated in this study is more susceptible to *M. marinum*. The increased infection burden of *cxcr3.3* mutants could be reverted to wild-type levels by injection of *cxcr3.3* mRNA, confirming the specificity of the mutant phenotype. A reduced infection burden, similar to the *cxcr3.2* mutant phenotype, was induced when *cxcr3.3* was overexpressed in wild-type AB/TL embryos, further supporting the notion that Cxcr3.2 and Cxcr3.3 have contrasting functions. We asked whether the underlying causes of the opposite effects of Cxcr3.2 and Cxcr3.3 on mycobacterial infection were due to essentially antagonistic functions or to a dysregulation of the transcription of the genes for the Cxcr3.2 and Cxcr3.3 receptors or the Cxcl11aa ligand. Gene expression profiles showed that *cxcr3.2* and *cxcl11aa* are induced upon infection with *M. marinum* and that *cxcr3.3* expression depends on *cxcr3.2*. The infection-driven induction of *cxcl11aa* remains unaltered in the *cxcr3.2* and *cxcr3.3* mutants, suggesting that the transcriptional regulation of the axis does not involve the ligand. While *cxcr3.3* expression levels were lower in *cxcr3.2* mutants, no alteration of *cxcr3.2* expression was detected in *cxcr3.3* mutants. Therefore, the increased infection susceptibility of *cxcr3.3* mutants cannot be explained by differences in the level of the functional Cxcr3.2 receptor or the Cxcl11aa ligand. Taking these data together, we propose that the Cxcr3-Cxcl11 signaling axis is regulated at least at two levels. At the transcriptional level, infection drives the expression of *cxcr3.2* (and indirectly *cxcr3.3*) and *cxcl11aa.* At the functional level, Cxcr3.2 signals in response to Cxcl1aa ligand, while Cxcr3.3, given its ACKR-like features, may function to negatively regulate Cxcr3.2 activity.

The increased infection burden of *cxcr3.3* mutants could either be due to a defective bacterial clearance or to altered macrophage migration properties, which can have major effects on the development of mycobacterial infection [29, 64, 8]. We demonstrated that *cxcr3.3* mutants can clear bacteria effectively and proceeded to evaluate if an altered macrophage migration could be facilitating bacterial dissemination. We obtained results supporting the functional antagonism between Cxcr3.2 and Cxcr3.3 when we locally injected *M. marinum* or purified Cxcl11aa protein into the hindbrain cavity. In both cases, we observed enhanced recruitment of macrophages to the site of injection in *cxcr3.3* mutants, while *cxcr3.2* mutants displayed reduced cell recruitment. Interestingly, while neutrophil recruitment was reduced in the *cxcr3.2* mutant, it remained unaltered in *cxcr3.3* mutants, suggesting that Cxcr3.3 has no effect on neutrophil migratory properties.

To examine whether altered cell motility was the underlying reason for enhanced recruitment in *cxcr3.3* mutant macrophages, we used a tail-amputation assay to assess migration in terms of total cell displacement and average speed. We showed that *cxcr3.3* mutant macrophages move faster than WT controls. To test our hypothesis, we assessed bacterial dissemination and confirmed that, in the context of *M. marinum* infection, the overall worse outcome in *cxcr3.3* mutant larvae was linked to amplified macrophage-mediated dissemination of bacteria that is facilitated by the higher speed of migrating macrophages and favors the formation of secondary granulomas. Since more macrophages were recruited when Cxcl11aa was injected into the hindbrain cavity and upon tail-amputation, we propose that the enhanced macrophage recruitment in *cxcr3.3* mutants is not a specific *M. marinum*-induced phenotype, but rather a Cxcl11-dependent response that can also result from wound-induced inflammation or other Cxcl11aa-inducing stimuli.

The circularity index (CI) is a measure indicative of the activation status of macrophages, with low CI values (stretched morphology) corresponding to a high activation status [65, 66]. The predominance of macrophages with low CI values in *cxcr3.3* mutants suggests that these cells have a higher activation status and that they are more responsive to stimuli in their environment. Cxcr3.3 depleted larvae showed an overall upregulation of inflammatory markers (*tnfa* and *cxcl11aa*) at 4 hpa. We suggest that the inflammatory phenotype of Cxcr3.3-deficient larvae reflects a dysregulation in the Cxcr3-Cxcl11 signaling axis, supported by the upregulation of *cxcl11aa*, that results in an exacerbated Cxcr3.2 signaling in the absence of the ligand-scavenging function of Cxcr3.3. In support of this model, the simultaneous chemical inhibition of the two Cxcr3 paralogs showed that only macrophages expressing Cxcr3.2 were affected and that the inhibitor treatment phenocopied *cxcr3.2* mutants regarding *M. marinum* burden and wound-induced macrophage recruitment. These data provide further evidence that Cxcr3.2 is directly involved in leukocyte trafficking, while Cxcr3.3 only fine-tunes the process by shaping the chemokine gradient and the availability of shared ligands.

Although we found that enhancement of Cxcr3.2 signaling due to the loss of Cxcr3.3 is detrimental in *M. marinum* infection, it might be beneficial in the context of other infections or in other processes dependent on macrophage recruitment, such as tissue repair and regeneration. Furthermore, it should be noted that the function of a chemokine receptor is primarily dependent on the type of cell expressing it, as the subset of receptors expressed by the cell and the intracellular integration of the signals have been shown to be determinant for functional specificity [28]. While our study revealed that macrophage migration is modulated by an antagonistic interplay between the Cxcr3.2 and Cxcr3.3 receptors, it remains to be determined how interactions between Cxcr3 paralogs may affect the function of other innate and adaptive immune cells. Although there is only one copy of CXCR3 in humans, there are 3 splice variants of the gene (CXCR3-A, CXCR3-B, and CXCR3-alt), and a regulatory mechanism for fine-tuning of CXCR3 function also exists. The splice variants CXCR3-A and CXCR3-B differ in their N and C termini and carry out antagonistic functions. CXCR3-A mediates chemotaxis and proliferation, while CXCR3-B inhibits cell migration and proliferation, and induces apoptosis [67, 68]. Both splice forms can bind to CXCL9-11 chemokines but mediate opposite functions. While there are no splice variants of *cxcr3.2* and *cxcr3.3* in zebrafish [69], the regulatory antagonism between the two paralogs resembles the interaction between the two human CXCR3 splice variants, which might suggest a form of convergent evolution. However, this functional diversification of CXCR3 variants is not conserved in the murine model, where CXCR3 is a single copy gene and no splice variants have been identified so far [30, 67].

In conclusion, our work illustrates the antagonistic interaction between the two CXCR3 paralogs Cxcr3.2 and Cxcr3.3 in zebrafish. The structural analysis of Cxcr3.3 supports that this receptor is unable to signal in the conventional G-protein-dependent way, but that it can still bind ligands and shape chemokine gradients, thereby regulating active receptors with shared ligands. Our experimental data show that the absence of the scavenging function of Cxcr3.3 is detrimental in the context of mycobacterial infection due to an exacerbated Cxcr3.2 signaling and a consequently enhanced macrophage motility that facilitates bacterial dissemination. However, we propose that enhanced macrophage motility could be benign in other contexts, such as tissue repair. Our findings suggest an extensive crosstalk between several chemokine signaling axes such as CXCR3-CXCL11 andCXCR4-ACKR3 (CXCR7), since ACKR3 also binds CXCL11 besides CXCL12 [26, 18]. Furthermore, ACKR1 a silent receptor that does not scavenge chemokines but redistributes them to mediate leukocyte extravasation, shares the CXCL11 and CXCL4 ligands with CXCR3 [70, 71]. The complexity of signaling axis integration, further emphasizes the relevance of unraveling the fundamental mechanistic principles underlying intricate chemokine networks. Our findings contribute to understanding one such mechanistic interaction and suggest that a more comprehensive ACKR classification needs to be developed to aid the understanding of complex chemokine signaling regulation.

## Abbreviations

ACKR: atypical chemokine receptor
CFU: colony forming units
CI: circularity index
dpf: days post fertilization
dpi: days post infection
EC: extracellular
GPCR: G-protein coupled receptor
hpa: hours post amputation
IC: intracellular
qPCR: quantitative PCR
TM: transmembrane
WT: wild-type

## Authorship

FS designed and performed experiments, analyzed the data, and wrote the manuscript. VT designed and performed experiments and analyzed data. SK and AL contributed to the experimental work. AM supervised the study and reviewed the manuscript. All authors commented on the manuscript and approved the final version.

## Acknowledgements

The authors thank Georges Lutfalla (University of Montpellier) and Steve Renshaw (University of Sheffield) for zebrafish reporter lines, Bjørn Koch for advice on time lapse imaging, and Gabriel Forn-Cuní for advice on the phylogenetic analyses. We are grateful to all members of the fish facility team for zebrafish care. FS was supported by a fellowship from CONACYT. VT was a Marie Curie fellow in the Initial Training Network FishForPharma (PITN-GA-2011-289209), funded by the 7^th^ Framework Programme of the European Commission.

