## Supplemental tables, figures and figure legends for "Antagonism between regular and atypical Cxcr3 receptors regulates macrophage migration during infection and injury in zebrafish"

**Supplementary materials**

**Supplementary table1.** Top hits for predicted ligands of Cxcr3.2 and Cxcr3.3.

| Cxcr3.2 | Ligand name | C-score |
|---|---|---|
| **1** | 0NN | 0.18 |
| **2** | DGW | 0.12 |
| **3** | 2CV | 0.5 |
| **4** | Y01 | 0.05 |
| Cxcr3.3 | Ligand name | C-score |
| **1** | DGW | 0.12 |
| **2** | 0NN | 0.10 |
| **3** | Y01 | 0.05 |
| **4** | 2CV | 0.04 |

**c-score-confidence index*

**Supplementary table 2**. Accession numbers of sequences used fir the phylogenetic tree.

| | Accession number |
|---|---|
| 1. Cavefish (CF) Ackr4 | ENST00000249887.2 |
| 2. Cavefish (CF) Ackr4b | ENSAMXG00000025769 |
| 3. Cavefish (CF)Cxcr3.2 | ENSAMXG00000035350 |

| | |
|---|---|
| 4. Cavefish (CF)Cxcr3.3 | ENSAMXG00000018866 |
| 5. Herring (CH) CXCR3 | XP_012694805.1 |
| 6. Cod (COD)Ackr3b | ENSGMOG00000019953 |
| 7. Cod (COD)Cxcr3.3 | ENSGMOG00000016951 |
| 8. Coelacanth (COE) Ackr2 | ENSLACG00000000319 |
| 9. Coelacanth (COE) Ackr3 | ENSLACG00000016030 |
| 10. Coelacanth (COE) Ackr4 | ENSLACG00000007877 |
| 11. Coelacanth (COE) Cxcr3.1 | XP_005999214.1 |
| 12. Coelacanth (COE) Cxcr3.2 | XP_014343707.1 |
| 13. Elephant shark (ES) Cxcr3 | XP_007909361.1 |
| 14. Frog (FR) Ackr3 | ENSXETG00000003296 |
| 15. Frog (FR) Cxcr3 | ENSXETG00000024989 |
| 16. Fugu (FU) Ackr4a | XP_003978861.2 |
| 17. Fugu (FU) Ackr4b | XP_011606262.1 |
| 18. Fugu (FU) Cxcr3.1 | XP_003966387.1 |
| 19. Fugu (FU) Cxcr3.3 | XP_003966388.2 |
| 20. Human (HU) Ackr1 | ENSG00000186810 |
| 21. Human (HU) Ackr2 | ENSG00000144648 |
| 22. Human (HU) Ackr3 | ENSG00000144476 |
| 23. Human (HU) Ackr4 | ENSG00000129048 |
| 24. Human (HU) Cxcr3 | ENSG00000186810 |
| 25. Lamprey (LAM) Ackr3 | XP_007909361.1 |
| 26. Mouse (MO) Ackr2 | ENSMUSG00000044534 |
| 27. Mouse (MO) Ackr3 | ENSMUSG00000044337 |

| | | |
|---|---|---|
| 28. | Mouse (MO) Ackr4 | ENSMUSG00000079355 |
| 29. | Mouse (MO) Cxcr3 | ENSMUSG00000050232 |
| 30. | Asian arowana (SF) Cxcr3 | XP_018587703.1 |
| 31. | Asian arowana (SF) Cxcr3a | KPP61297.1 |
| 32. | Spotted gar (SG) Ackr2 | ENSLOCG00000009409 |
| 33. | Spotted gar (SG) Ackr4a | ENSLOCG00000018134 |
| 34. | Spotted gar (SG) Ackr3 | ENSLOCG00000004865 |
| 35. | Spotted gar (SG) Cxcr3.1 | XP_015224255.1 |
| 36. | Spotted gar (SG) Cxcr3.3 | AWO96327.1 |
| 37. | Zebrafish (ZF) Ackr3a | ENSDART00000090414 |
| 38. | Zebrafish (ZF) Ackr3.2 | ENSDARG00000058179 |
| 39. | Zebrafish (ZF) Ackr4a | ENSDART00000145446 |
| 40. | Zebrafish (ZF) Ackr4b | ENSDART00000058703 |
| 41. | Zebrafish (ZF) Cxcr3.1 | ENSDARG00000078177 |
| 42. | Zebrafish (ZF) Cxcr3.2 | ENSDARG00000041041 |
| 43. | Zebrafish (ZF) Cxcr3.3 | ENSDART00000146611 |

**Supplementary figure 1. Neutrophil development is unaltered in *cxcr3.3* mutants.**

No evident morphological aberrations were observed in *cxcr3.3-/-* larvae within the first 5 dpf and the numbers of neutrophils in whole body (A), head (B), and tail (C) were unaltered in *cxcr3.3* and *cxcr3.2* mutants. The cell numbers corresponding to each day are the average of 35 larvae of each of the 3 groups (genotypes). Data were analyzed using a two-way ANOVA and are shown as mean ± SEM (ns $p > 0.05$, * $p \leq 0.05$, ** $p \leq 0.01$, *** $p \leq 0.001$, **** $p \leq 0.0001$).

**Supplementary figure 2. Injection of an empty CMV vector had no effect on bacterial burden after systemic *Mm* infection.**

WT and *cxcr3.3* mutant larvae were injected with an empty CMV vector at 0 hpf and bacterial burden was assessed at 4dpi. The injection of the empty vector had no effect on the outcome of *Mm* infection on either group (A). We conducted a One-way ANOVA to test for significance. Results are plotted as mean ± SEM (ns $p > 0.05$, * $p \leq 0.05$, ** $p \leq 0.01$, *** $p \leq 0.001$).

**Supplementary figure 3. *dram1* mutant larvae cannot clear mycobacterial infection efficiently.**

*dram1* mutant embryos and their WT siblings were systemically infected at 28 hpf with 200 CFU of the *ΔERP* wasabi *M. marinum* strain. The total number of bacterial clusters in every fish was quantified **(A)** and bacterial clusters were divided into 3 groups based on the number of bacteria they contained (1-5,1-6 and > 10) to assess bacterial clearance at 44 hpi **(B).** Dram1-depleted larvae had more bacterial cluster than their WT siblings and developed larger bacterial clusters.

**Supplementary figure 4. Cxcr3.3 mutants have a predominantly pro-inflammatory phenotype.**

To assess the activation status of WT, *cxcr3.2* and *cxcr3.3* mutant larvae, we assess the transcriptional profile of the three M1 markers, *tnfa* **(A),** *cxcl11aa* **(B)** and *Il1b* **(C)** at 4hpa. In *cxcr3.3* mutants the former two (A-B) were upregulated while there was no difference in il1b (**C**) among the three groups

**Online supplementary videos 1.** Representative time-lapses of WT, *cxcr3.2* mutants, and *cxcr3.3* mutant larvae after tail amputation. Time-lapses show macrophages of WT, *cxcr3.2* mutants, and *cxcr3.3* mutant larvae migrating towards the injury. Images from a z-stack of the injured area  were acquired every 60 sec for 1.5 h and combined into max projection time-lapse.
